# Profiling cellular diversity in sponges informs animal cell type and nervous system evolution

**DOI:** 10.1101/758276

**Authors:** Jacob M. Musser, Klaske J. Schippers, Michael Nickel, Giulia Mizzon, Andrea B. Kohn, Constantin Pape, Jörg U. Hammel, Florian Wolf, Cong Liang, Ana Hernández-Plaza, Kaia Achim, Nicole L. Schieber, Warren R. Francis, Sergio Vargas R., Svenja Kling, Maike Renkert, Roberto Feuda, Imre Gaspar, Pawel Burkhardt, Peer Bork, Martin Beck, Anna Kreshuk, Gert Wörheide, Jaime Huerta-Cepas, Yannick Schwab, Leonid L. Moroz, Detlev Arendt

## Abstract

The evolutionary origin of metazoan cell types such as neurons, muscles, digestive, and immune cells, remains unsolved. Using whole-body single-cell RNA sequencing in a sponge, an animal without nervous system and musculature, we identify 18 distinct cell types comprising four major families. This includes nitric-oxide sensitive contractile cells, digestive cells active in macropinocytosis, and a family of amoeboid-neuroid cells involved in innate immunity. We uncover ‘presynaptic’ genes in an amoeboid-neuroid cell type, and ‘postsynaptic’ genes in digestive choanocytes, suggesting asymmetric and targeted communication. Corroborating this, long neurite-like extensions from neuroid cells directly contact and enwrap choanocyte microvillar collars. Our data indicate a link between neuroid and immune functions in sponges, and suggest that a primordial neuro-immune system cleared intruders and controlled ciliary beating for feeding.

---

Sponges are sister to all, or nearly all other animals (Fig. 1A) (*1, 2*). Unlike most animals, they lack bona fide neurons, muscles, and a gut. Rather, their body plan is composed of three basic cell types that act to create an efficient water pump for filter-feeding and waste removal (Fig. 1, B to C, and fig. S1). First, the choanocytes form spherical chambers, and exhibit microvilli and a cilium that beats to drive water through the canal system (Fig 1C and fig S1, J to O). The pinacocytes are epithelial cells that line the canals and form the outer cover and the basal layer adherent to the substratum. Lastly, mesenchymal cell types fill the inner space (mesohyl) between the choanocyte chambers and the pinacocytes, and include stem and skeletogenic cells.

**Figure 1.**
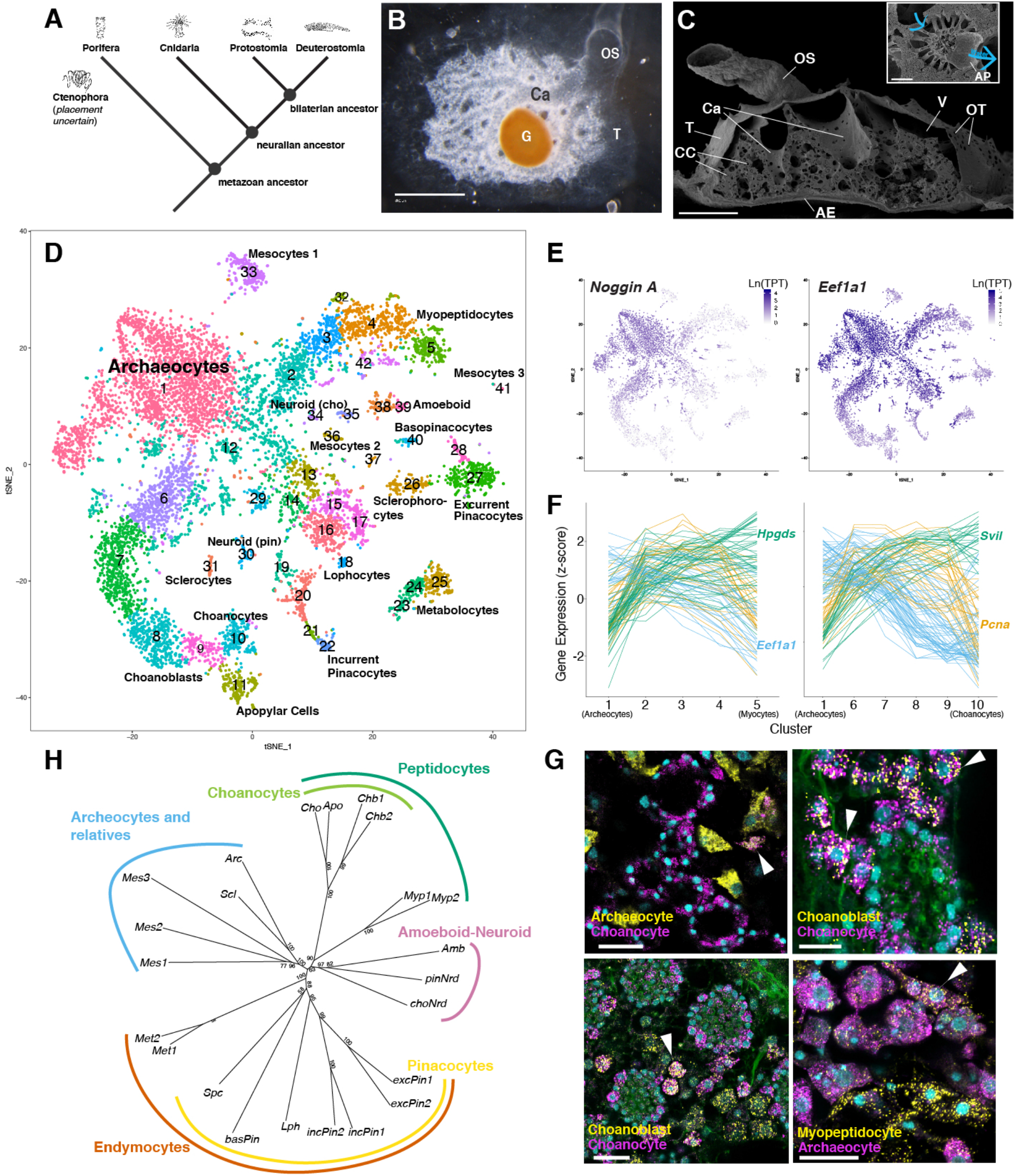
*Spongilla lacustris* cell types from whole-body scRNAseq. (A) Simplified phylogeny of major animal groups (with sponges as sister to all other animals). (B) Overhead view of juvenile *S. lacustris*. Scale bar 300 μm. (G) gemmule, (Ca) excurrent canals, (T) outer epithelial tent, (Os) osculum. (C) SEM showing lateral cross-section of juvenile *S. lacustris*. Water enters through ostia (OT) into the vestibule (V) and is drawn through the canal system via choanocyte chambers (CC; inset), before exiting through the osculum. Scale bar 300 μm. (AE) attachment epithelium. (D) tSNE plot of 10,106 individual cells colored according to 42 genetically-distinct clusters. (E) Archaeocyte markers show gradual reduction in expression along cell trajectories. (F) Expression along myocyte (left) and choanocyte (right) trajectories of the top 70 differentially expressed genes for clusters 2 and 3 (left) and clusters 6 and 7 (right). Genes colored by expression pattern across trajectory: increasing (green), decreasing (blue), increasing followed by decreasing (orange). (G) smFISH of archaeocyte (*Eef1a1*), myopeptidocyte (*Hpgds*), and choanocyte (*Villin-like*), and choanoblast (*Pcna*) markers showing differentiated and transitional (arrow) cells with observed co-localization of selected gene markers. Cyan – nuclei (DAPI), green – membrane (Cellbrite Fix). Scale 20 μm (10 μm in top right panel). (H) Neighbor-joining tree of Spongilla cell types. Node numbers indicate bootstrap support (10,000 replicates).

Despite their simple organization, sponges posess genes encoding conserved molecular machinery found in neurons, muscles and additional specialized cell types of bilaterian animals. This includes components of the bilaterian neuronal pre- and postsynapse (*3*). In addition, sponges show various forms of slow movements, including rhythmic contractions that flush the canal system and expel debris (*4*). However, cells with integrative signaling functions in sponges are unknown, and the genetic signature and relationships of most cell types are incomplete. Sponges are therefore a key lineage for inferring the early history of animal cell type evolution and the origins of the nervous system.

## Sponges are composed of 18 distinct cell types

Using whole-body single-cell RNAseq, we conducted a comprehensive survey of genetically distinguishable cell types in *Spongilla lacustris*, a freshwater Demosponge. We captured and barcoded mRNA from individual cells of dissociated 8-day old *S. lacustris* juveniles, a stage at which they have acquired all mature morphological features, including numerous choanocyte chambers, well-developed canal system, and differentiated mesohyl. Mapping was performed against a *de novo* transcriptome, for which we inferred high-quality gene annotations using an automated phylogenetic pipeline that provided detailed orthology information for every *Spongilla* gene (*5*). In total, we obtained expression data from 4 single-cell capture experiments, resulting in a dataset of 10,106 individual *S. lacustris* cells, expressing 26,157 genes out of 39,552 in the *S. lacustris* transcriptome (fig. S2, A to C).

Using Louvain graph clustering, we resolved 42 cell clusters with distinct gene expression signatures. Projected into 2-dimensional expression space, this revealed a central cluster from which other clusters emanated (Fig. 1D and fig. S2D). Marker genes for this cluster (Data S1) included *Noggin, Musashi1*, and *Piwi-like*, that demarcate the pluripotent, stem celllike archeaocytes (*6, 7*). Expression of archaeocyte markers gradually decreased with distance from the central cluster to the peripheral clusters (Fig. 1E), and clusters interconnecting the center and periphery exhibited expression profiles intermediate between those of the archaeocytes and peripheral clusters (Fig. 1F), with few unique markers (Data S1). In contrast, clusters in the periphery showed much more unique expression profiles, comprised of both transcription factors and effector genes (fig. S2, E to F, and Data S1), indicating the presence of distinct differentiation programs for these cells. Based on this, we interpreted the “arms” emanating from the center as developmental trajectories, extending from archaeocyte stem cells via committed precursors towards differentiated cells in the periphery. The peripheral clusters accordingly represent differentiated cell types of the sponge. In four cases, the trajectory endpoints comprised a pair of closely related clusters, with hundreds of genes specific for the cluster pair, but relatively few genes specific to each individual cluster, suggesting these pairs represent distinct states of the same cell type (fig. S2, E to I).

We next selected marker genes for each cluster and visualized their expression using single-molecule fluorescent *in situ* hybridization (smFISH; Data S1). This resolved spatial relationships of stem cells, developmental precursors, and differentiated cell types across the entire organism. Probes directed against archaeocyte markers labeled a population of large mesenchymal cells with a prominent nucleolus (Fig. 1G and fig. S4, A to C). Peripheral clusters 10 and 11 were identified as choanocytes and apopylar cells based on expression of the actin-binding protein *Villin* (Fig. 1G), along with other known choanocyte markers (*8*). Clusters 8 and 9, which fall intermediate between archaeocytes and choanocytes, exhibited upregulation of many proliferation markers, including *Pcna*. We interpreted these as choanoblasts, in agreement with previous reports that choanocyte precursors are highly proliferative (*9*). Confirming this, we found that cells coexpressing *Vilin* with *Pcna* or the archaeocyte marker *Eef1a1* often neighbored choanocyte chambers (Fig. 1G). Following similar strategies, we assigned all peripheral clusters to a total of 18 differentiated cell types (Table S1), representing a novel unbiased classification of cell populations in Porifera. Concordant with previous morphological description, we recovered pinacocytes and several previously-described mesenchymal cell types, such as the sclerocytes, which produce the siliceous spicules, and cells that transport spicules (*10*). However, other mesenchymal cell types in our dataset did not match any previous classification and thus represent newly discovered cell types.

## Hierarchical organization of 4 cell type families

Cell types in most animals are organized into distinct families, which share expression of regulatory and effector genes, and have been identified in recent studies using hierarchical clustering (*11*). To explore cell type interrelationships in sponges, we first used weighted correlation network analysis (WGCNA) to identify gene sets that co-varied across differentiated cell types (fig. S3A). This revealed 28 distinct gene sets, some of which delineated unique cell types, whereas others represented groups of cell types, hinting at the presence of hierarchical organization. To test this explicitly, we applied a statistical model that assessed whether cell type interrelationships based on gene expression have more hierarchical organization than expected by chance. This assigns a “treeness” score for every combination of four cell types (a tetrad), and estimates a p-value using randomized datasets (fig. S3B). We found strong support for hierarchical structure in gene expression patterns among cell types, with close to 60% of cell type tetrads exhibiting greater hierarchy than expected by chance (fig. S3, C to D). This result was consistent regardless of whether using all expressed genes or only variable genes, or whether using log-normalized counts per million (CPM) or binarized “on-off” expression values (fig. S3C).

Based on this, we generated a cell type tree using neighbor-joining tree reconstruction (Fig. 1H), revealing four well-supported clades of differentiated cell types. This included the extended pinacocyte and choanocyte families, as well two families of primarily mesenchymal cells, one containing immune and neuroid cells, and another with sclerocytes and enigmatic mesocyte cell types (fig. S4). Notably, there is no clear separation between epithelial and mesenchymal cells in our tree. Rather, we observed mesenchymal cell types scattered across all four cell type clades, including well-supported relationships between choanocytes and myopeptidocytes, and between the epithelial pinacocytes and mesenchymal lophocytes and sclerophorocytes.

## Endymocytes are contractile and respond to nitric oxide signaling

We refer to the family of pinacocyte-related cell types as endymocytes (from greek ἔvδνμα: “lining, clothing”), which have in common to cover and shape the sponge body. Among endymocytes, the incurrent pinacocytes constitute both layers of the tent, the lining of the vestibule, and the outer layer of the osculum (Fig. 2, A to D, and fig. S5, A to D). Excurrent pinacocytes line the excurrent canals and inner osculum layer (Fig. 2E and fig. S5, D to E), shaping the entire excurrent system. Lastly among epithelial endymocytes, the basopinacocytes comprise the basal epithelial layer attached to the substrate (Fig. 2G and fig. S5, F to G). Mesenchymal members of the endymocyte family, including collagen-secreting lophocytes (Fig. 2H and fig. S5H), sclerophorocytes (Fig. 2I and fig. S5I), and metabolocytes (Fig. 2J and fig. S5, J to K), were observed most abundantly near the basal layer and in the periphery of the sponge, often in proximity to pinacocytes.

**Figure 2.**
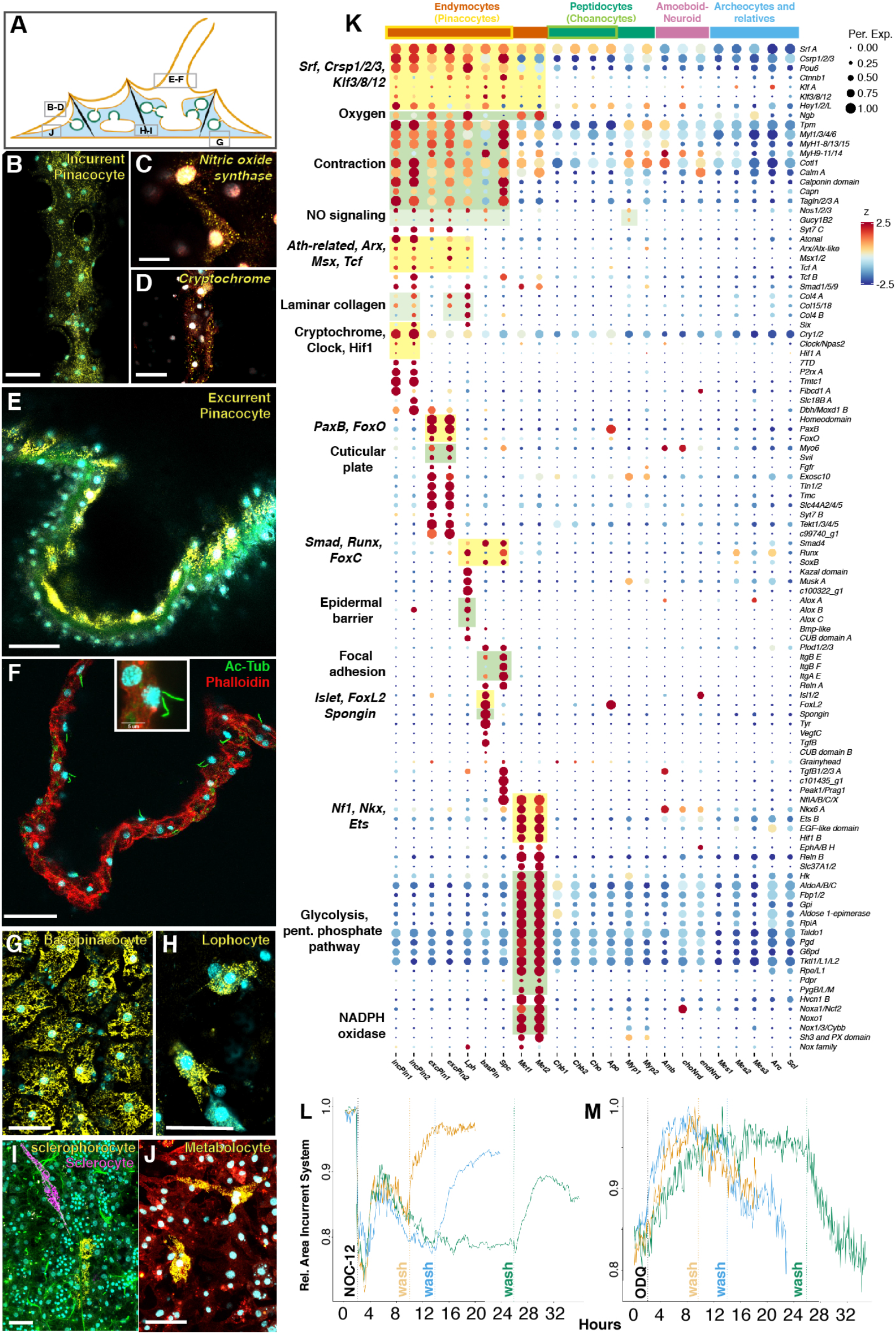
Endymocyte cell type family. (A) Sponge diagram illustrating locations of corresponding smFISH panels for endymocyte cell types. Modified from Schippers and Nichols (*57*). (B–J) smFISH of peptidocyte markers. Markers: (B) *Tropomyosin*, (E) *c99740_g1*, (G) *Spongin short-chain collagen*, (H) *c100322_g1*, (I) *Silicatein* (sclerocyte) and *c101435_g1* (sclerophorocyte), (J) *Reelin*. Membrane stains Fm-143Fx (red) and CellBrite Fix (green; also apparently stains collagen fibrils); cyan – nuclei (DAPI); Scale bar 30 μm. (K) Dotplot of endymocyte cell type markers. Boxes around dots delineate expression modules of transcription factors (yellow) or functional complexes/pathways (green). (L-M) Timelapse plot of contraction state following NOC-12 (C) or ODQ (D) treatment. Colors indicate experiments with treatment for 8 (yellow), 12 (blue), or 24 hours (green).

Gene ontology analysis of endymocyte gene modules found strong enrichment for Wnt and TGF-β signaling (fig. S5L), as well as a whole actomyosin-contractility module (Fig. 2K), consistent with findings in the marine demosponge *Amphimedon queenslandica* (*12*), and with pinacocytes being the main contractile cell types mediating whole-body rhythmic contractions in sponges (*13*). We also observed expression of the transcription factors *Serum response factor (Srf)*, a master regulator of the cellular contractile apparatus (*14*), and *Cysteine and glycine-rich protein (Csrp1/2/3*, aka *Muscle LIM protein*) encoding cofactors of Srf, suggesting sponges may contain a conserved regulatory module for actomyosin contractility (Fig. 2K).

In other demosponges, signaling molecules such as GABA, glutamate, and nitric oxide have been shown to alter rhythmic contractions (*13, 15, 16*). Supporting this, we found paralogs of the metabotropic GABAB receptor specifically employed in distinct endymocytes (fig. S6). The pinacocytes also specifically co-expressed *Nitric oxide synthase*, which catalyzes the synthesis of nitric oxide (NO), as well as a putative NO receptor, *guanylate cyclase 1 soluble beta 2* (*Gucy1B2*) (Fig. 2K). NO triggers relaxation of vertebrate endothelial muscle (*17*), and has been implicated in regulating sponge contractions (*13, 16*). Based on this, we sought to characterize contractions in *S. lacustris* and the role of NO in regulating this behavior. We first developed a general model of contractions in *S. lacustris*. An individual contraction cycle in *S. lacustris* occurs over seven distinct stages, each driven by coordinated morphological changes in incurrent, excurrent, and oscular tissue systems (fig. S7, A to B, and Movie S1). During the first three stages, contraction of the osculum and incurrent systems cause water to swell excurrent canals (fig. S7A). In stages 4 and 5, the excurrent system contracts, expelling water through the osculum. In the final stages, both incurrent and excurrent systems return to the resting state (fig. S7A).

We next tested the response of juvenile *Spongilla* to NO by treating juvenile sponges with NOC-12, a small molecule donor of NO, and also ODQ, an inhibitor of the NO receptor *Gucy1b2* (Fig. 2, L to M, fig. S7C, and Movies S2-3). Addition of NO resulted in an immediate strong contraction of the incurrent system, including canals, ostia and tent (Fig. 2L), along with the simultaneous expansion of the excurrent canal system, similar to that observed during the initial stages of an endogenous contraction. In the continued presence of nitric oxide, the incurrent system remained in a contracted state, undergoing a series of short, pulsed contractions. Full relaxation occurred only after the removal of nitric oxide by washing, which also resulted in the return of both incurrent and excurrent systems to resting states (Fig. 2L). The effect of ODQ treatment was subtler, although we observed relaxation of incurrent, and possibly excurrent, canal systems upon treatment (Fig. 2M), which was only relieved by washing. We also conducted experiments in which NOC-12 and ODQ were added in serial combination, which revealed generally similar results, with NOC-12 inducing strong contractions, and ODQ causing relaxation (Fig. S7C). These results show nitric oxide plays a key role in both initiating and regulating contractions in *S. lacustris*, and possibly all demosponges (*13*).

Beyond contraction, cell type-specific expression programs indicated specialized roles for endymocytes in sensation, skeleton formation, metabolism, and defense. Incurrent and excurrent pinacocytes express single orthologs of *Six* and *Pax*, the two main homeodomain factors of the conserved Six/Pax regulatory network (*18*), in a complementary manner (Fig. 2K). *Six+* incurrent pinacocytes express the blue-light sensor *Cryptochrome* (Fig. 2K), suggesting these cells are light-sensitive. *PaxB+* excurrent pinacocytes express other sensory markers, including the mechano-electrical transducer protein *Tmc*, as well as *Supervillin* and *MyosinVI* (Fig. 2K), which encode proteins that regulate the rigid F-actin meshwork in sensory hair cells. Validating this, we observed short primary cilia on excurrent pinacocytes in the osculum and excurrent atrium (Fig. 2F), in line with previous reports (*19, 20*). In addition, we found basopinacocytes enwrapping spicules anchored at the base of the sponge (fig. S5, F to G), and sclerophorocytes forming clusters alongside mature spicules (Fig. 2I and fig. S5I), reminiscent of previously described *SoxB+* spicule “transport cells” in *E. muelleri* (*10*). In line with skeletogenic functions, basopinacocytes and sclerophorocytes express several integrin paralogs, known to connect cells to the basal lamina via focal adhesions (Fig. 2K). The most distinct endymocyte family member are the newly discovered metabolocytes, large multipolar mesenchymal cells with wide extensions that may contact pinacocytes (Fig. 2K and fig. S5, J to K). Metabolocytes specifically upregulate the complete set of enzymes mediating glycolysis and the pentose phosphate pathway (Fig. 2K), and uniquely express *glycogen phosphorylase* and *pyruvate dehydrogenase*, indicating they are key regulators of energy metabolism in the sponge. Metabolocytes specifically express *nox* genes encoding subunits of NADPH oxidase (Fig. 2K), a membrane-bound enzyme complex that faces the extracellular space and produces reactive oxygen species to kill intruding bacteria during phagocytosis (*21*).

## Peptidocytes are digestive and express receptive ‘post-synaptic’ scaffolding genes

We refer to the choanocyte-related cell type family as peptidocytes (from greek πέπτειν: “to digest”), reflecting their shared role in digestion. Peptidocytes comprise the choanocytes proper and a closely related cell type that we identified as apopylar cells (*Apo*) using smFISH (Fig. 3, A to C). Apopylar cells form the excurrent pore of choanocyte chambers (Fig. 3A and fig. S1M-N), and are thought to regulate water currents inside of the sponge via control of the diameter of the efflux opening (*22*). We also discovered myopeptidocytes, an abundant yet previously undescribed constituent of the mesohyl (Fig. 3, D to E). These form long projections that contact each other, and other cells including choanocytes. Uniquely among peptidocytes, they express many contractile genes, including three paralogs of *Mib2* (Fig. 3F), known to regulate myofiber integrity (*23*). The presence of mesenchymal cells expressing the contractile apparatus is striking, as contraction in sponges has been shown to primarily involve pinacocytes (*24, 25*), although a role for the mesohyl had been suggested (*4*).

**Figure 3.**
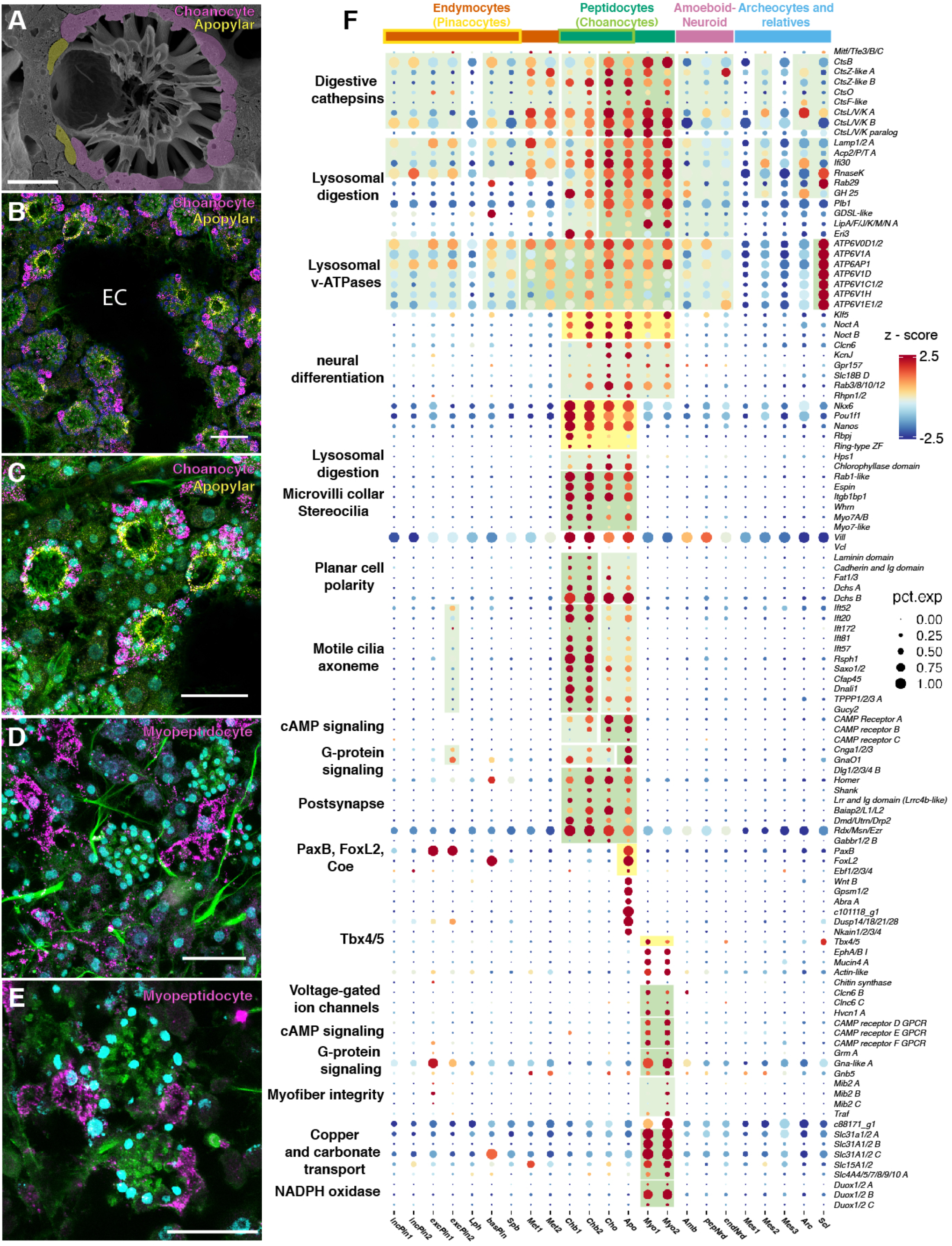
Peptidocyte cell type family. (A) SEM of choanocyte chamber showing choanocytes (purple) and apopylar cells (gold). Scale bar 10 μm. (B – E) smFISH of peptidocyte markers. Markers: (B-C) *Villin-like* (choanocytes) and *c101118_g1* (apopylar cells), (D-E) *c88171_g1*. Membrane stain CellBrite Fix (green) also apparently stains collagen fibrils; cyan – nuclei (DAPI); EC - excurrent canal. Scale bar 30 μm. (F) Dotplot of peptidocyte cell type markers. Boxes around dots delineate expression modules of transcription factors (yellow) or functional complexes/pathways (green).

Recent volume EM studies reveal that the choanocyte internal volume is primarily composed of food vacuoles (*26*), making digestion a pivotal function. Corroborating this, all peptidocytes are strongly enriched for lysosomal gene sets (Fig. 3F), including lysosomal biogenesis markers and digestive enzymes such as cathepsins (Fig. 3F). Choanocytes also express the transcription factors *Rbpj, Klf5*, and *NK homeobox 6* (Data S2 and S3), known to regulate digestive cell types, endodermal development, and metabolism in other animals (*27–29*). Notaby, we also observed expression of six members of the *Sorting nexin* family with Phox and BAR domains (*SNX-PX-BAR*) (Fig. 3F) involved in macropinocytosis, an unregulated form of engulfment (*30*). This is consistent with choanocytes engulfing non-food particles, such as latex beads, and bringing them into lysosomes (*31*).

Significantly, we observed highly specific expression of the core postsynaptic density proteins *dlg, homer*, and *shank* (Fig. 3F) in the choanocyte subclade – and a broader survey revealed additional choanocyte-specific ‘postsynaptic’ genes, involved in membrane organization and linking transmembrane signaling components to the actin cytoskeleton, such as *Dystrophin/Utrophin* (Fig. 3F and fig. S6) (*32*). Notably, several of these genes, including the sole ortholog of *Ezrin, Radixin*, and *Moesin*, as well as *Baiap2* and *Eps8* also play an important role in organizing the microvilli actin skeleton. This suggests postsynapse-like scaffolding machinery may in fact be localized to choanocyte microvilli.

## The amoeboid – neuroid family shows an innate immunity profile

This family comprises three small cell types dispersed in the mesohyl. They lack a nucleolus, and are triangular or multi-angular in shape with long, filopodia-like extensions (Fig. 4, A to F, and fig. S8, A to D). Whereas the choano-neuroid cells associate with choanocyte chambers (Fig. 4, A to D, and fig. S8, A to B), the amoeboid and pinaco-neuroid cells are dispersed in the mesohyl (Fig. 4, C to F, and fig. S8, C to D). We also observed pinaco-neuroid cells embedded next to pinacocytes, including incurrent pinacocytes of the outer tent (Fig. 4E and fig. S8D).

**Figure 4.**
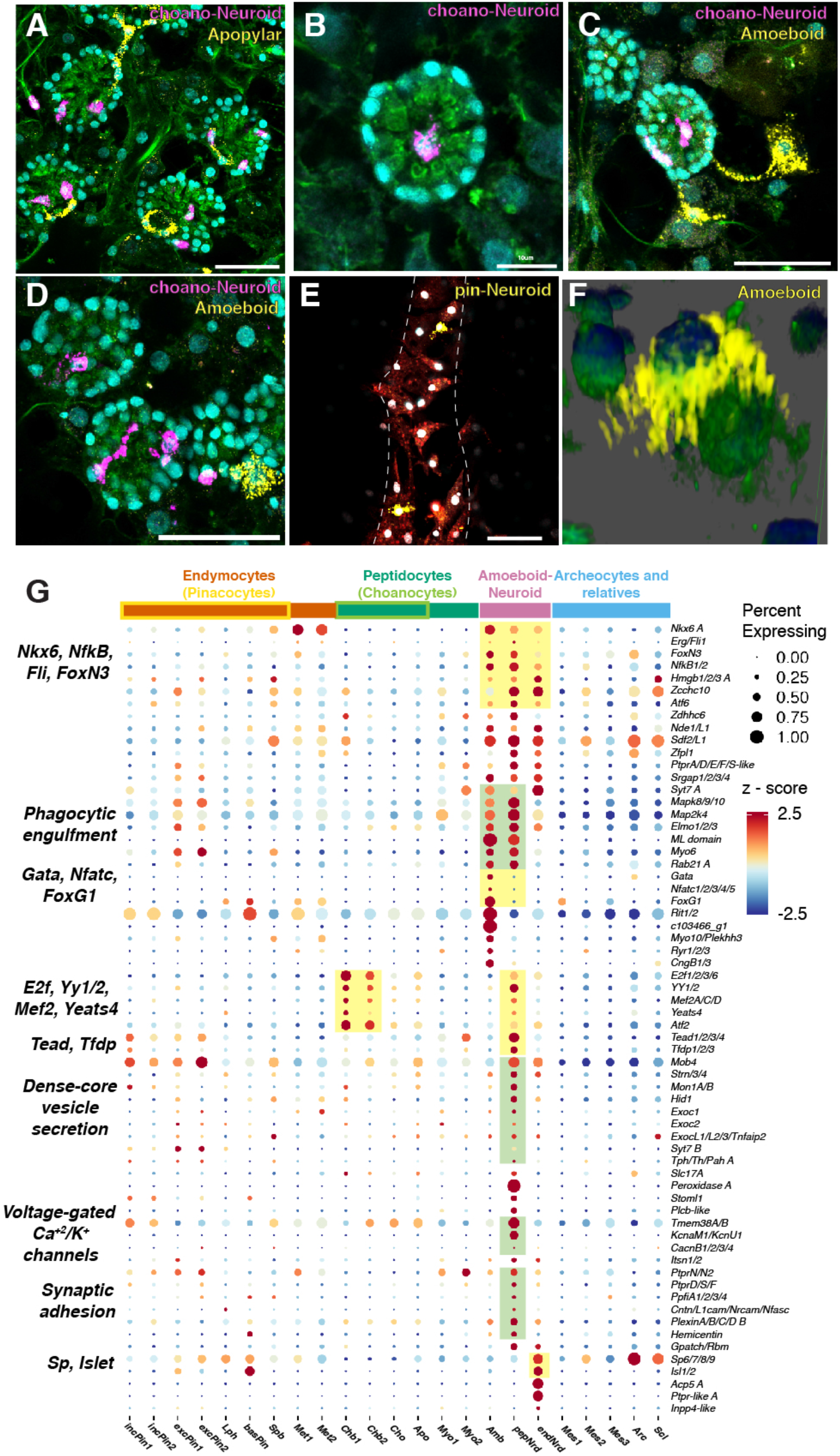
Amoeboid-neuroid cell type family. (A-F) smFISH of amoeboid-neuroid markers. Markers: *Peroxidase A* (choano-Neuroid), *c85989_g1* (Apopylar cells), *c103466_g1* (Amoeboid cells), *Acp5* (pinaco-Neuroid). Dotted line outlines epithelial tent. Membrane stains are CellBrite Fix (green) and Fm143-Fx (red); cyan – nuclei (DAPI). Scale bar 30 μm unless noted in the panel. (G) Dotplot of neuroid-amoeboid cell type markers. Colored boxes delineate transcription factor (yellow) or effector (green) modules.

Family-specific transcription factors suggest a role in innate immunity. *Nuclear factor kappa B subunit (NFkB*) controls cellular immune responses, and *High mobility group box 1 (Hmgb1*) (Fig. 4G and fig. S2F) acts as both secreted cytokine and nucleic-acid sensor, in addition to its nuclear role, activating key inflammatory pathways important for innate immunity (*33*). Further, amoebocytes specifically express the only sponge orthologs of *Nfat*, which plays important roles in both adaptive and innate immunity, and *Gata*, an evolutionarily conserved determinant of immunocytes (*34*).

Family-specific effector genes support phagocytic function and the formation of filopodia (Fig. 4G). This includes the evolutionarily conserved *engulfment and cell motility (Elmo*) (*35*), essential for engulfment of dying cells in worms and phagocytosis in mammalian cells (*35*), the small GTPase *Rab21*, fundamental for phagosome formation (*36*), and *Myosin-X (Myo10*), known to induce filopodia (*37*). Validating this, we repeatedly observed amoeboid cells engulfing other cells in the sponge mesohyl (Fig. 4F), suggesting a role cell corpse removal. Engulfment and filopodia formation have in common to rely on targeted exocytosis, which supplies the excess membrane required for filopodia extension, or for the formation of the engulfing ‘phagocytotic cup’. In line with this, the amoeboid-neuroid cell types are enriched for genes involved in the generation and sorting of vesicles, polarized vesicle trafficking, and targeted exocytosis of vesicles (Fig. 4G).

## Communication between neuroid, choanocytes, and apopylar cells

Beyond innate immunity, we observed a unique ‘presynaptic’ expression module for the choano-neuroid cells, indicating specific interaction and communication with the ‘postsynaptic’ choanocytes. In vertebrates, LAR family receptor-type protein tyrosine phosphatases (RPTPs) are crucial for synapse formation on the presynaptic side (*38*), interacting with the Ptprf-interacting protein a (*Ppfia*; aka liprin-α) that links to the presynaptic scaffold. Strikingly, we found the sponge LAR *PtprD/S/F* and *Ppfia1/2/3/4* uniquely expressed in choano-neuroid cells, as was the single sponge ortholog of *PtprN* protein tyrosine phosphatases important for dense-core vesicle secretion (*39*) (Fig. 4G). Beyond this, choano-neuroid cells were enriched for genes involved in neurosecretory trafficking and dense core vesicle secretion (Fig. 4G). They also expressed more general ‘presynaptic’ genes mediating vesicle priming and SNARE exocytosis, although these genes were also expressed in other cell types, including many endymocytes (fig. S6). On the postsynaptic side, vertebrate RPTPs interact with different binding partners, such as the Leucine-rich repeat and Ig domain-containing protein Netrin-G ligand 3 (*38*). In the sponge, we found a Leucine-rich repeat and Ig-like domain-containing protein similar to netrin-g3 ligand (*Lrrc4b*) uniquely expressed in choanocytes and apopylar cells (fig. S6E). This suggests that LAR, Liprin-α, and RPTP-binding protein mediate interaction between choano-neuroid and choanocytes/apopylar cells, in line with the presence of postsynaptic-scaffold-like matrix in the latter.

To challenge this hypothesis, we tested for spatial contact between choano-neuroid, choanocytes, and apopylar cells. Inspecting smFISH and confocal microscopy images, we noted that nearly all choano-neuroid cells were associated with choanocyte chambers, either forming part of the choanocyte lining, or located in the middle of chambers (Fig. 4, A to D, and fig. S8). This was reminiscent of the sponge ‘central cells’, which have been described to populate choanocyte chambers in some demosponges, although not in *S. lacustris*, and to contact surrounding choanocytes with long cellular protrusions (*40*). Indeed, we observed long cellular extensions protruding from choano-neuroid cells lining the chambers, as well as from neuroid cells more central, with cellular protrusions contacting individual choanocytes (Fig. 4, A to D).

We then used correlative light electron microscopy to find back choano-neuroid cells in a 30μm^3^ FIB-SEM volume (Fig. 5). Using machine-learning, we segmented the entire volume and reconstructed the spatial relationships and interactions of all cells within a choanocyte chamber, identifying two likely choano-neuroid cells situated inside the chamber (Fig. 5, A to F). The first choano-neuroid cell was positioned in the center of the chamber (violet cell; Fig. 5A), with the second cell residing near the apopylar pore (red cell; Fig. 5B). Our reconstruction revealed that both neuroid cells form multiple protrusions, each directed towards the collar of individual choanocytes (Fig. 5, B to F). Strikingly, nearly all extensions from the choano-Neuroid cells contact and enwrap one or more microvilli (Fig. 5, D to E), with each extension reaching to a different choanocyte collar. In several cases, we also observed choano-Neuroid cell extensions in close proximity to cilia, and even orienting themselves along the main ciliary axis. Cilia in close contact with the neuroid protrusions emerged straight, and then bent, contrasting with the undulatory, corkscrew-like appearance of normal motile cilia, suggesting they may not be actively beating.

**Figure 5.**
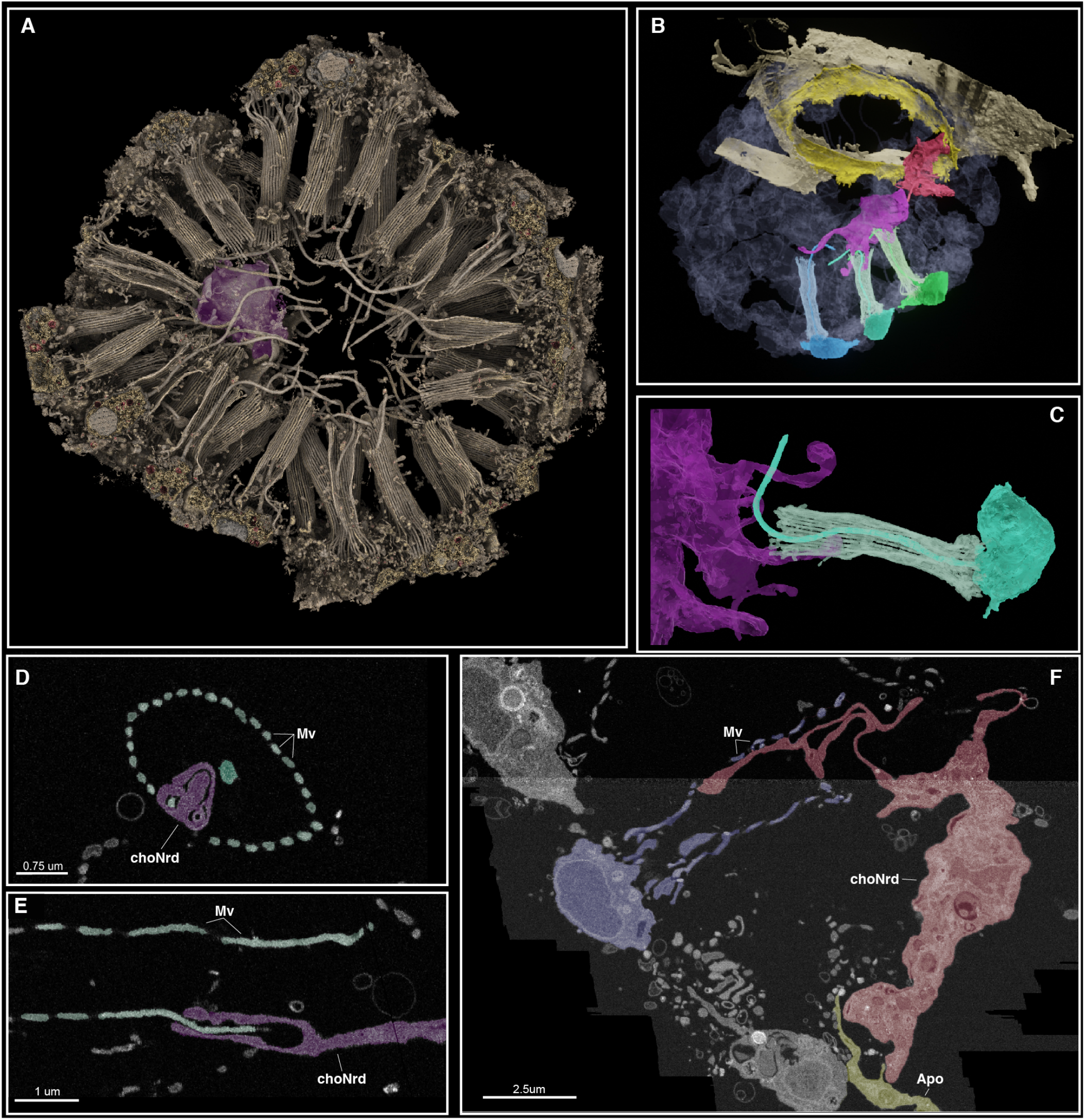
FIB-SEM of neuroid-choanocyte interaction. (A) Rendered 3D volume of choanocyte chamber with choano-Neuroid cell (violet). (B) Segmented volume showing two choano-Neuroid cells (violet and red) contacting cilia and microvillar collars of three choanocytes (blue, turquoise, and green) and apopylar cells (yellow). (C) Segmented choano-Neuroid cell (violet) with filopodia extending into the microvillar collar (turquoise). (D-F) 2D images of choano-Neuroid cell filopodia (purple and red) extending into, and enwrapping, choanocyte microvilli (Mv; turquoise and dark blue), and contacting apopylar cell (Apo; yellow).

## DISCUSSION

A notable feature of gene and species evolution is the prevalence of a tree-like history of common descent, the first evidence for which was based on Darwin’s observation that traits among living organisms are organized hierarchically. In our study, we observed significant hierarchical organization of gene expression among all differentiated cell types of a sponge, which was reflected in the combinations of transcription factors and functional modules shared by different subclades in the *S. lacustris* cell type tree. We propose that such shared expression among differentiated cell types reflects the history of cell type evolution (*41*). Under this model of cell type evolution, the major sponge cell type families - pinacocytes, choanocytes, neuroid/amoeboid, and sclerocyte relatives – originated from ancestral cell types that further diversified within sponges. Such evolutionary lineages are distinct from developmental trajectories, which are instead constructed based on the gradual change and overlapping expression of developmental genes controlling the transition from multipotent precursor to differentiated cells (*42*).

For peptidocytes, the expression of cathepsins and lysosomal enzymes, biogenesis genes, and vATPases, suggests they principally function as digestive cells. Peptidocytes also specifically express a Klf ortholog most similar to *Klf5*, which play a conserved role in digestive epithelia specification (*28, 43*), and choanocytes express *Nkx6, Rbpj*, and *Notch*, the vertebrate orthologs of which specify endocrine and exocrine cell types in pancreas (*44*) and absorptive enterocytes (*27, 45*). *Nkx6* also demarcates pharyngeal and exocrine gland cells of the digestive filaments in the sea anemone *Nematostella* (*46*). Klf5+, Nk6+, Rbpj+ peptidocytes thus likely existed in metazoan ancestors, which may have initially fed via intracellular digestion – as observed in sponges and in the unicellular relatives of animals (*26, 47*) – and then acquired the capacity to target digestive enzymes into exocrine vesicles for external digestion in other animals – first as part of a digestive mucoid-ciliary sole, and then later incorporated into the gut (*48*). In line with an evolutionary transition from intra- to extracellular digestion, conventional lysosomes can also be subject to exocytosis (e.g. in pancreatic acinar cells) (*49*).

In endymocytes, our experiments reveal complex coordination between sensory-contractile pinacocytes, coordinated in part through nitric oxide signaling, which is also known to regulate endothelial smooth muscle contraction in vertebrates (*17*). The pinacocyte transcription factor signature supports a possible evolutionary link between sensory-contractile systems in sponges and in other animals. In both *Spongilla* (our data) and in the calcisponge *Sycon ciliatum* (*50*), pinacocytes specifically express the sole sponge ortholog of *Msx*, which plays a conserved role in specifying sensory ectodermal regions across animals (*51*), but was originally named after its function in myocyte specification (*52*). *Msx* is known to activate *atonal*, which plays a conserved role in hair cell and mechanosensory neuron specification in various animals (*53*), and is specifically expressed in sponge pinacocytes, together with its bHLH heterodimeric binding partner *Tcf4/E12/47*. Most revealing, excurrent pinacocytes and incurrent pinacocytes express the single sponge orthologs of the *Pax* and *Six* homeodomain transcription factors, respectively. In other animals, *Pax* and *Six* paralogs play conserved roles in the specification of sensory cells (e.g. 54), but others have been implicated in myocyte specification (*55*). Indeed, in bilaterians, an entire conserved network comprising *Pax, Six, Eya* and *Dach* genes functions in both the specification of sensory organs and musculature and it is intriguing that in sponges both functions may relate to the same clade of sensory-contractile cells. Taken together, our data are consistent with the notion that sensory cells and myocytes arose by the division of labor from shared evolutionary precursors, and that these precursors existed as sensory-contractile, conducting epithelia in metazoan ancestors.

With amoeboid-neuroid cells, the specific expression of key immunity markers including *NFkB, Hmgb1, NFAT*and *GATA* indicate an affinity to immune cells, or hemocytes, present in virtually all other animals (*34*). In addition, however, our data also indicate a role for choano-neuroid cells in intercellular communication, and ultimately liken them to neural cell types of other animals (Fig. 6). The close physical interaction of neuroid filopodia with choanocyte microvilli suggests a coordinative role in feeding, by modulating ciliary beating. *Spongilla* is a sponge with open architecture, where incoming particles between 5μm and 50μm collect on the outer choanocyte chamber walls and must be continuously removed to prevent clogging (*40*). The neuroid cells may take part in this process by ‘turning off’ the cilia of individual chambers for surface cleaning. Together, our data are consistent with recent observations that nerve nets play a role in the tuning of defense mechanisms in holobiont organisms (*56*), and suggest that immune and nervous systems evolved by the division of labor from an initial neuro-immune system – a derived variant of which is found in extant sponges. Such a system might have captured and/or monitored bacteria, and at the same time regulated microbial food uptake in response to quality and availability.

**Figure 6.**
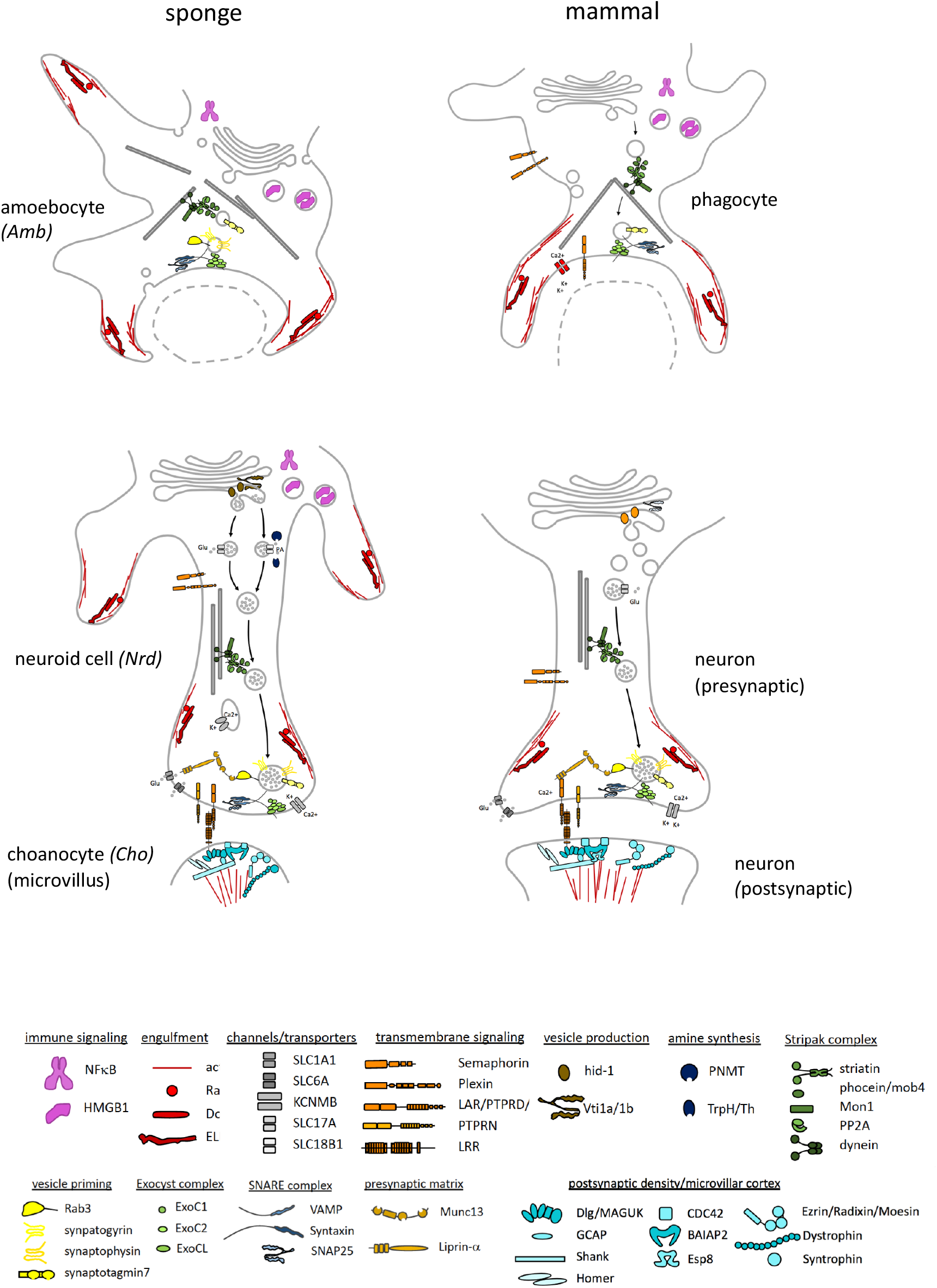
Sponge and mammalian cell types share expression of cellular modules with dual neural/immune functionality. Shared proteins active in phagocytic cup formation (sponge amoebocyte and mammalian macrophage) and targeted vesicle release (sponge choano-neuroid cell and neuron). The cellular location of sponge proteins is hypothetical.

## Acknowledgements

We thank the Advanced Light Microscopy Facility and the Electron Microscopy Core Facility at the EMBL for support.

## Author contributions

Conceptualization, J.M.M., M.N., L.L.M., D.A.; Methodology, J.M.M, M.N., A.B.K., Y.S., L.L.M., D.A.; Software, J.M.M., M.N., C.P., C.L., A.H-P., W.R.F., J.H-C.; Validation, J.M.M, K.J.S., S.K., M.R.; Formal Analysis, J.M.M., M.N., C.P., C.L., A.H-P., W.R.F, J.H-C.; Investigation, J.M.M., K.J.S., M.N., G.M., A.B.K., J.U.H, F.W., K.A., N.L.S., S.K., M.R., I.G., Writing - Original Draft, J.M.M., D.A.; Writing – Review & Editing, J.M.M., K.J.S., M.N., L.L.M, D.A.; Visualization, J.M.M., K.J.S., M.N., G.M., F.W., A.H-P., D.A.; Supervision, J.M.M, M.N., P.Bu., P.Bo., M.B., A.K., G.W., J.H-C., Y.S., L.L.M., D.A.; Funding Acquisition, L.L.M., D.A.

