## Supplementary Materials and Methods for "Profiling cellular diversity in sponges informs animal cell type and nervous system evolution"

**This PDF file includes:**

Materials and Methods
Figs. S1 to S8
Table S1
Captions for Movies S1 to S5
Captions for Data S1 to S3

**Other Supplementary Materials for this manuscript include the following:**

Movies S1 to S5
Data S1 to S3.

### Materials and Methods

#### Collection and cultivation of sponges

During winter, *Spongilla lacustris* individuals form protected packets of stem cells called gemmules, which can be collected and stored in the lab, and induced to hatch and form juvenile sponges. Adult specimens of *S. lacustris* in overwintering stage were collected on 12 January 2017 and 8 March 2018 from Lake Constance, near Kressbronn, Germany (@ 47°35'09.0"N 9°35'56.4"E). Individual sponge patches, composed of the sponge skeleton and gemmules, were stored in lake water at 4°C until gemmule isolation. Gemmules were extracted from the adult tissue via gentle agitation of the skeleton structure by lightly rubbing the sponge over sandpaper (grit 500) using one finger covered in nitrile gloves. Gemmules from different sponge patches were washed several times with either Vittel mineral water or M-medium (1 mM $CaCl_2 \cdot 6H_2O$, 0.5 mM $MgSO_4 \cdot 7H_2O$, 0.5 mM $NaHCO_3$, 0.05 mM KCl, 0.25 mM $Na_2SiO_3$) (*58*) and stored separately at 4°C in Vittel mineral water.

Juvenile sponges were grown by placing gemmules in culture dishes with M-medium, and kept at room temperature (20-25°C) in the dark. To culture sponges for 10x single-cell RNAseq and smFISH, 4-8 gemmules were placed in a glass-bottom culture dish (Greiner Bio-One cat# 627860) containing 5 ml of Vittel mineral water. To culture sponges for Immunostaining, 4-8 gemmules were placed on a coverslip (24x24 mm, #1) inside 6-well plates, each well containing 5 ml of M-medium. To culture sponges for the contraction experiments, ~4 gemmules were placed on a coverslip (24x32 mm, #1) inside a 60 mm culture dish containing 10 ml of M-medium.

M-medium was replaced every other day, beginning around day 5 when juvenile sponges first adhere to the cover slip. Juvenile sponges were sampled or fixed for experiments after 7-12 days of growth, when they had acquired all major features of adults, including an osculum, well-developed canal system, and numerous choanocyte chambers. Variance in the exact time at which this stage occurs is extremely high, even for neighboring sponges grown from clonal gemmules that are present in the same culture dish. In general we tended to err on the side of letting sponges develop further, so that all sponges in the dish exhibited these features.

#### Construction and annotation of transcriptome

**Bulk RNA isolation and sequencing**

RNA was extracted from juvenile *S. lacustris* using Trizol (Thermofisher #15596026) and the resulting extracts were quantified using a Nanodrop 2100 and quality controlled on a Bioanalyzer 2100. We conducted 100PE and 125PE sequencing on a HiSeq 1500 using an RNA extract with RIN 10, generating 10,459,248 and 12,198,962 read pairs for the two samples, respectively.

**Trinity de-novo transcriptome**

Bulk RNA-seq reads were adapter trimmed, quality filtered and assembled using Trinity (*59*), with the in-silico normalization option selected and other parameters set as default. This resulted in a transcriptome N50 of 905 base pairs. A BUSCO (*60*) search using the metazoa dataset indicated that the reference transcriptome was fairly complete (95.2% BUSCOs found). We identified putative proteins using Transdecoder (version 3.0.1), requiring a minimum open reading frame length of 100 amino acids.

**Phylome and transcriptome refinement**

To infer orthology relationships between genes in the *S. lacustris* de-novo transcriptome with genes of other animals, we generated a comprehensive phylome containing trees for all genes in the *Spongilla* transcriptome. Briefly, each inferred *S. lacustris* protein was used as a seed to construct a gene/protein tree. For each seed, identification of the 250 closest homologs in 53 proteomes from other metazoan and unicellular relatives was performed using BLAST with e-value and query coverage cutoffs of $E10^{-3}$ and 33% respectively. Phylogenetic workflow was executed using the workflow template named "clustalo_default-trimal001-none-raxml_default" from the ETE Toolkit v3.1.1 (*61*), which consisted of multiple sequence alignment reconstruction using clustalOmega v1.2.1 (*62*) with default parameters, Trimal (*63*) for alignment cleaning by removing columns with more than 1% gaps, and Maximum Likelihood tree inference using RAXML v8.1.20 (*64*) with default parameters.

All the resulting trees were rooted to the farthest leaf to each seed sequence using the function 'get_farthest_oldest_leaf' from the ETE Toolkit and using the NCBI Taxonomy tree as a reference to established relative dating of the species included. Rooted trees were automatically processed using ETE's `get_descendant_evol_events` function to infer speciation and duplication nodes. Speciation nodes were used to infer fine-grained orthology relationships (one-to-one, one-to-many, many-to-many) between seed sequences and other species.

We next refined the assignment of transcripts to genes in our de-novo transcriptome using information derived from our phylome. We identified *S. lacustris* genes whose transcripts and proteins did not align to each other, but which mapped as one-to-one orthologs to the same gene in the closely related freshwater demosponge *Ephydatia muelleri*. Visual inspection of many individual cases suggested these *S. lacustris* Trinity genes were fragments (e.g. 5' and 3') of the same gene. In a few cases, these also represented genes that were incorrectly fused in the *Ephydatia muelleri* transcriptome, although this represented only a small minority of cases (<5%). Based on this, we grouped transcripts that did not align to each other, but were mapped as one-to-one orthologs to the same *E. muelleri* protein, into a single gene. This final assignment of transcripts to gene identifiers was used to construct a GTF file for mapping our single-cell 3' RNAseq data and quantifying UMI counts for each gene.

**Functional annotation**

We assigned orthology information and gene ontology terms to each gene in our final refined transcriptome. Preliminary gene names, used in our GTF annotation file and uploaded single-cell dataset, were assigned based on their ortholog relationship to human genes in our phylome. For some *S. lacustris* genes, we were not able to identify a clear human ortholog(s). For these we assigned names by mapping the longest protein for a gene to orthology groups in the eggNOG database (version 4.5.1) using eggNOG-mapper v1 (*65*). We inferred secreted peptides using a previously established pipeline (Moroz et al., 2014). Lastly, we assigned gene ontology terms for each Spongilla gene using eggNOG-mapper v1.

For all gene names used in the manuscript, we assigned a final name after visual inspection of the gene tree, eggnog mapper result, and domain structure. Gene names in the manuscript follow a standardized nomenclature, with names based on the official human names of all orthologs. For instance, a sponge gene orthologous to human genes *Lamp1* and *Lamp2* is named Lamp1/2. In cases where there are multiple sponge genes that are co-orthologs of the same human gene(s), these are given alphabetic designators (e.g. *Lamp1/2 A* and *Lamp1/2 B*). Lastly, in cases where the gene is inferred to be a paralog of a human gene (i.e. no human ortholog exists), the gene is appended with "like" (e.g. *Myo7-like*).

**Homeobox and Klf gene trees**

Individual phylogenetic trees were reconstructed for Homeobox and KLF gene families using a more exhaustive phylogenetic pipeline. First, a cluster of homologous sequences for each gene family was manually selected using BLAST searches based on e-value and score to include all relevant transcripts and known genes from reference species. For the Homeobox gene family, all sequences with score >= 100 were included. For KLF, 647 sequences with at least 75 score were selected. The software UPP (*66*), specifically designed to improve the alignment of short fragments, was then used with parameters -M1 -B100 to infer a multiple sequence alignment for each family. Finally, a phylogenetic tree was inferred using IQTree with parameters -m TEST -bb 1000 -bnni

#### 10x single-cell 3' RNAseq

**Sample collection and sequencing**

Dissociation of 8-day old juvenile *S. lacustris* in tissue culture dishes was performed by replacing M-medium with calcium and magnesium-free dPBS. Sponges were gently agitated by repeated pipetting, causing relatively rapid dissociation of all cells. Once completely dissociated (~5-10 minutes), cells were passed through a 40 µm mesh filter twice and collected in a low-bind 1.5 ml Eppendorf tube. Cells were washed twice by spinning at 1000g followed by resuspension in a small volume of calcium and magnesium-free dPBS. Cell viability was assessed using a combination of propidium idiodide (Cat #P4170, Sigma) and fluorescein diacetate (Cat #F7378, Sigma) to ensure greater than 90% live cells were used for capture. We conducted 4 single-cell RNAseq capture experiments from 8-day old juvenile *S. lacustris* using the 10x Chromium Controller and Chromium Single Cell 3' Kit v2 (Cat #120237, 10x Genomics, USA).

cDNA synthesis and library construction were made according to manufacturer's recommendations. Post-library quality control was determined on the Qubit fluorometer (Cat# Q32866, ThermoFisher Scientific) with the Qubit dsDNA HS Assay Kit and 1:10 sample dilution was ran on the Agilent 4200 TapeStation system (Cat#G2964AA, Agilent Technologies) with the High Sensitivity D1000 ScreenTape (Cat#5067-5584, Agilent Technologies) and High Sensitivity D1000 Reagents (Cat#5067-5585, Agilent Technologies). Post-library quantification was performed with Illumina Library Quantification Kit (Cat #KK4824, KAPA Biosystems).

Single-cell libraries were sequenced on an Illumina NextSeq500 using a 2x150 paired-end kits with the following sequencing read recommendation Number of Cycles: 26 cycles Read1 for cell barcode and UMI, 8 cycles I7 index for sample index and 98 cycles Read 2 for the transcript.

**Demultiplexing and mapping 10x data**

Individual 10x sample libraries were demultiplexed using Cell Ranger Makefastq v2.1.1 with default settings. Reads for each sample were mapped and demultiplexed by cell barcode and UMI using Cell Ranger Count v2.1.1. Lastly, we constructed a final aggregated count matrix using Cell Ranger Aggregate, which subsampled reads to ensure equal sequencing depth between all four samples.

**QC, clustering, and gene set analysis**

We visually inspected the distribution of genes, UMIs, and % mitochondral genes across cells. Ultimately, we chose to use a relatively inclusive threshold, removing cells expressing fewer than 200 genes, resulting in a final count matrix of 10,106 cells. We also tested other criteria for removing cells, but did not find that more stringent cutoffs yielded significantly different clustering results.

We explored a variety of clustering methods, ultimately using a combination of high-resolution Louvain clustering, implemented in R v3.5.1 using Seurat v2.4 (*67*), combined with a custom script for merging similar clusters (see GitLab repository). For this, we first selected higly variable genes using the approach of Macosko et al. (*68*). The top ~1000 genes were then used to perform principal components analysis. To identify principal components with significant variation, we performed a jackstraw resampling test, visually inspected the elbowplot of principal component vs. principal component standard deviation, and examine the genes with high loading for each principal component. Based on this we selected the top 40 principal components for cell clustering. However, we also tested using as few as 20, and as most as 100, of the top principal components, which generally yielded similar clustering results.

Clustering was conducted using the Louvain algorithm in Seurat. We explored extensively how Louvain clustering worked under different parameters, and on different datasets (e.g. stricter thresholds for cell inclusion, data subsets, etc). In general, clustering results were highly robust to different clustering parameters, with the exception of the resolution parameter, which significantly affected how many clusters we recovered. We found that clustering with higher resolutions was necessary to recover several rare but distinct cell types. However, this resolution setting also subdivided more common cell clusters, such as archaeocytes, into many smaller clusters that did not exhibit unique marker genes.

To identify all distinct clusters in our dataset, we opted for high-resolution clustering followed by iteratively merging nearest-neighbor clusters based on a similarity threshold. Briefly, clustering was first performed with the resolution parameter set to 10. We then calculated average expression profiles for each cluster, and used normalized expression vectors to calculate pearson correlations between each pairwise cluster combination. These were then ranked from highest to lowest correlation, and a Wilcoxon rank sum test was used to calculate the number of differentially expressed genes between each pair. Cluster pairs which differed by less than 20 differentially expressed genes with 2 fold change were merged. This process was performed iteratively until all clusters were sufficiently distinct. Using this approach, we identified 42 genetically distinct clusters in our dataset.

We used the number of “true” markers to distinguish between clusters representing putative cell types from those representing developmental transition states. “True” markers for a cluster were defined as those markers that were most highly expressed in that cluster, as inferred using the Wilcoxon rank sum test. Clusters representing transitional states were designated as those clusters with less than 30 "true markers", whereas those with more were designated as putative differentiated cell types. In total, we identified 23 putative cell types and 19 transitional states. Ultimately, we linked these 23 genetically distinct cell types to the 18 distinct morphological types referred to in the main text (with several morphological types exhibit multiple semi-distinct expression states). Lastly, we visually inspected the position of putative cell types and transitional clusters by generating a tSNE plot with default parameters in Seurat. Inspection of transitional clusters confirmed they occupied expression states intermediate between endpoints on the trajectories in our tSNE plot.

We identified co-expressed gene sets using weighted correlation network analysis, implemented in the R package WGCNA v1.68 (*69*). For this, we first identified an expanded set of ~6000 variable genes in Seurat to use in the analysis. We ran WGCNA on this pool of genes using mostly default parameters. Using the recommended approach, we set the softPower parameter to 5, required a minimum module size of 20, and conducted a final merging of gene sets whose eigengenes exhibited greater than 90% correlation.

**Hierarchy test and cell type tree**

To evaluate the level of tree structure among genetically distinct cell types in our single cell RNA-seq dataset, we first calculated the average value of log normalized counts per ten thousand for each putative cell type. We then calculated two types of distance matrices among cell clusters: the Euclidean distance for continuous expression values, and the Hamming distance for discretized values. We used a threshold of 0.025 for discretizing expression values, such that around 40% of the genes are marked as expressed in our analysis. Next, we followed the method in Liang, et. al. (*70*) to calculate the level of tree structure, $\pi_0$. Specifically, we calculated the $\delta$-value for each group of four cell clusters (a tetrad) directly from the distance matrix. In total we estimated $\delta$ for a total of 8855 tetrads. $\delta$ values close to 0 indicate that the distance matrix of the tetrad conforms the requirement of a tree, whereas $\delta$ values close to 1 indicate that the tetrad lacks tree structure. We then estimate the significance of $\delta$ that comes from non-random gene expression using the formula: $p = \frac{3\sqrt{3}}{2\pi(\delta^2 - \delta + 1)}$. In this dataset, we obtained 23 different cell clusters and a total of 8855 tetrads. The Benjamini and Hochberg (BH) multiple test correction implemented in the 'qvalue' R package was utilized to estimate the fraction of non-random tree structure, $\pi_0$, which serves as an indicator of the level of tree structure in the dataset. Finally, the distance matrix was visualized using the SplitTree program (*71*).

To construct the *Spongilla* cell type tree, we calculated a distance matrix with the 'dist' function in R using average log normalized expression profiles for each of our 23 genetically distinct cell types, and restricting genes to the same ~6000 variable genes used in our WCNA gene set analysis. Neighbor-joining tree reconstruction was performed with the R package 'ape' v5.3. To test support for each node in the tree, we performed 10,000 bootstrap replicates. The final tree was visualized using the Interative Tree of Life (iTOL) version 3 (*72*).

#### Single-molecule FISH

Prior to fixation, juvenile sponges growing in glass-bottom culture dishes were incubated with 200 µl of CellBrite Fix 488 membrane staining solution (stock diluted 1:1000 in Vittel mineral water or M-medium; Cat #30090, Biotium) for 15 min at RT, washed twice with Vittel water, and then fixed with 2% paraformaldehyde in ¼ Holtfreter's solution (¼ HS: 87.5 mg NaCl, 1.25 mg KCl, 2.5 mg CaCl2, 5 mg NaHCO3 in 100 mL $H_20$, Funayama et al. 2005) for 20 min at RT (room temperature). Specimens were washed once in ¼ HS for 5 min, then dehydrated with 50% MetOH in ¼ HS, 100% MetOH, and 100% EtOH. Specimens were then permeabilized with 200 µl of 50% Xylene/EtOH for 15 min at RT, while trying to minimize the exposure of Xylene to the plastic sides of the dish as much as possible. Specimens were rehydrated in a series of 75%, 50% and 25% EtOH in PBS-Tr (PBS with 0.1% Triton-X100), and then washed three times in PBS-Tr for 10 min each. Sponge specimens were transferred to 1 ml Hybridization buffer (2x SSC pH 7.0, 15% Ethylene Carbonate, 1 mM EDTA, 50 µg/ml heparin, and 1% Triton X-100 and were pre-hybridized for at least 10 min at 37°C. Hybridization was carried out overnight at 37°C with 3 nM labelled ssDNA oligos (i.e. smFISH probes) in 200 µl Hybridization buffer. smFISH probes were constructed against target RNA following Gaspar et al. (*73*). DNA Oligos (Data S1) were bound to either ddUTP-Atto633 (Cat #25040, Lumiprobe and Cat #AD633, Atto Tec) or ddUTP-Atto565 (Cat #Ad565, Atto Tec) fluorophores. During hybridization, the dishes were covered with parafilm to prevent evaporation. After hybridization, samples were washed with 1 ml Hybridization buffer (without smFISH probe) three times for 15 min each at 37°C. Specimens then were washed three times with 3 ml PBS at RT. For DNA counterstaining, sponge samples were incubated with 1 ml DAPI staining solution (1 ug/ml DAPI in PBS; Cat #D9542, Sigma-Aldrich) for 20 min at RT and subsequently washed twice with 3 ml PBS. Sponge specimens were stored in PBS at 4°C for up to one week. Images were taken with a 40x water immersion objective on a Leica SP8 inverted confocal microscope, and images were processed using Huygens deconvolution software and ImageJ (z-stack: maximal projection intensity projection).

#### Immunostaining

Immunostaining procedure was performed as described previously (*57*). Primary antibody incubation was with Acetylated tubulin (1:500 dilution; Cat #T6793, Sigma-Aldrich), and secondary antibody incubation was with Alexa Fluor 488 Goat, Anti-Rabbit IgG (H+L; 1:500 dilution; Cat #111-545-144, Jackson ImmunoResearch), Alexa Fluor 568 Phalloidin (1:40 dilution; Cat #A12380, ThermoFisher), and DAPI (1 ug/ml). Images were taken with a 63x oil immersion objective on a Leica SP8 inverted confocal microscope, and images were processed using ImageJ (z-stack: maximal projection intensity projection) and Imaris x64 v9.3.1.

#### Sponge Contraction Experiments

##### Imaging endogenous contractions

Time-lapse imaging for the general contraction model was performed as modified from Nickel (*25*) using a computer controlled Nikon Coolpix 990 attached to a dissecting microscope (Zeiss SM XX or Zeiss Stemi 2000) through a custom made projection adapter (Zeiss GF-PK 10/20 ocular glued to a Kendo 28 mm adapter ring). The time-lapse was triggered through a PC running DC-Time-Trigger and DC-Remote-Shutter 2.0.1. software (Per Madsen, digitalkamera.dk). Illumination was achieved by cold-light source (Zeiss KL 200) set on low intensity in order to avoid heating up the sponges.

Sponges were imaged in 60 mm petri dishes. For bi-planar views, a coverslip (24x24 mm, Roth) was gold-sputtered in a sputter coater (Emitech K500) to serve as a mirror, which was introduced at a 45° angle as close as possible to the imaged sponge. The mirror coverslip was precisely positioned using a 3-way micro-manipulator (Zeiss). The camera field of view was adjusted through the dissecting microscope zoom lens to fit both views simultaneously, the axial view as well as the mirror-based lateral view.

**Image processing of endogenous contractions**

Image analysis and time-lapse video creation was performed in ImageJ. For measurements in axial and lateral views, grey value thresholds were used to represent total projected area of lateral and axial projected areas, based on a custom made loop macro, measuring a whole image stack. Projected areas of subdermal lacunae, incurrent canals, excurrent canals and oscule were manually measured at given positions (oscule in lateral view) or partial areas (M1-M3 in axial and lateral views).

**Flow-through chamber for NO experiments**

We utilized a flow-through system to enable introducing and flushing away substances to juvenile sponges without directly touching the dish or chamber holding the sponge. This is critical, as mechanical vibration is known to stimulate sponge contractions. Juvenile sponges grown on coverslips were inserted into a micro observation chamber (*74*) of 1.7 ml volume which was placed on an inverted microscope (Zeiss Cell Observer). The chamber was part of a custom made gravity driven continuous flow system, with reservoir containing fresh M-medium placed on a shelf above the microscope, and connected to the chamber via flexible medical plastic tubing. A collection reservoir, also connected to the chamber via medical tubing, was placed below the chamber. Partial closure of the tubing enabled for precise control of the flow-rate through the chamber containing the sponges, which was typically set between 10-50 mL per hour. We built in luer lock adapters and 3-way-valves to allow for the introduction of experimental substances via injection with a 30 mL syringe.

**Nitric oxide and ODQ experiments**

We introduced NOC-12 (Cat #E3145, Sigma-Aldrich) or ODQ (Cat #O3636, Sigma-Aldrich) separately and in serial combination to juvenile sponges in our flow-through chamber. We also performed control experiments in which we halted flow-rate or introduced DMSO. For each experiment, juvenile sponges were introduced into the flow system 4-12 hours before injection, allowing them to equilibrate under a constant flow rate of 10-50 mL M-medium per hour. Following this, we gently injected 15 mL of the treatment solution, which ensured that the observation chamber and all tubing leading in an out of chamber, up to the valves, were flushed with the treatment solution. For experiments with single treatment solutions, we alternated incubation periods with washing periods in which normal flow rates of M-medium were resumed. For experiments containing both NOC-12 and ODQ, we first incubated sponges with one substance, followed by incubation with both substances, before lastly washing by resuming normal flow of M-medium.

| **Experiment** | **Treatment solution** | **Concentration** | **Experimental sequence** |
|---|---|---|---|
| Nitric oxide | NOC-12 | 22.7 µM | Normal flow (12 hrs), NOC-12 (12 hrs), wash/normal flow (12 hrs), NOC-12 (24 hrs), wash/normal flow (12 hrs), NOC-12 (8 hrs), wash/normal flow (48 hrs) |
| sGC inhibition | ODQ | 3 µM | Normal flow (26 hrs), ODQ (8 hrs), wash/normal flow (13 hrs), ODQ (12 hrs), wash/normal flow (12 hrs), ODQ (24 hrs), wash/normal flow (48 hrs) |
| Nitric oxide followed by sGC inhibition | Noc-12, ODQ | 22.7 µM, 3 µM | Normal flow (21 hrs), NOC-12 (8 hrs), NOC-12/ODQ (16 hrs), wash/normal flow (12 hrs), NOC-12 (12 hrs), NOC-12/ODQ (24 hrs), wash/normal flow (11 hrs), NOC-12 (24 hrs), NOC-12/ODQ (16 hrs), wash/normal flow (48 hrs) |

| | | | |
|---|---|---|---|
| sGC inhibition followed by Nitric oxide | ODQ, Noc-12 | 3 µM, 22.7 µM | Normal flow (19 hrs), ODQ (2 hrs), ODQ/NOC-12 (11 hrs), wash/normal flow (11 hrs), ODQ (9 hrs), ODQ/NOC-12 (16 hrs), wash/normal flow (12 hrs), ODQ (17 hrs), ODQ/NOC-12 (27 hrs), wash/normal flow (48 hrs) |
| Control | DMSO | | Normal flow (24 hrs), DMSO (8 hrs), wash/normal flow (12 hrs), DMSO (24 hrs), wash/normal flow (24 hrs) |

**Time-lapse imaging of nitric oxide experiments**

During contraction experiments, images were taken using a consumer camera (Panasonic G2), attached to the microscope through a custom made ocular adapter through a MFT-T2 adapter which included an 8x projective (Olympus). The camera was connected to a flash unit (Nikon SB24), covered by a red diffuser foil. Both, camera and flash were connected to a permanent power source (Panasonic DMW-BLB13 and custom made). Time lapse imaging was triggered by a intervalometer set either to 30 or 60 sec interval. Images were stored on the camera using a wifi-SDcard (ez Share), which was connected to a PC running ez Share Windows Client V1.1.0 under Microsoft Windows, transferring Images directly to a network drive after capturing.

**Image processing of treated sponges**

Images were processed in FIJI in order to remove background noise on the coverslip surrounding the sponge, which tends to accumulate over time. A semiautomated macro was used to segment the total sponge area: Based on the wand tool functionality the macro was provided with a pixel coordinate inside the sponge, plus a lower (depending on the single experiment, which slightly varied due to the sponges in use) and upper grey (255) value threshold, from which the threshold area was grown based on 8-connectivity. The resulting outline of the sponge was expanded by 10 pixels in all directions and the background removed (set to 0/black). This was automated across the time stacks taken.

Segmenting images of whole sponges is challenging as they exhibit a complex three dimensional topology. To infer relative size of mesohyl, choanoderm, and canal compartments, we used grey value thresholding (*15*), tuning thresholds independently for each sponge to best capture each compartment. Relative pixel measurements were obtained for the full time series in FIJI and plotted in R using 'ggplot2' v3.1.1.

#### FIB-SEM of Neuroid – Choanocyte Interaction

**Pre-fixation and Fluorescence Confocal Microscopy:**

*Spongilla* were grown on Ibidi Glass Bottom Dishes 35 mm for 14 days and were prefixed with aldehydes and stained with DAPI. Due to the high osmotic sensitivity of these freshwater animals, we performed the pre-fixation in two steps: first for 30 min at room temperature in 4% FA in ¼ HS medium and subsequently in 2.5% GA and 1.6% FA in PHEM in a microwave (Pelco Biowave containing a ColdSpot cooling system, 150-W on/off cycling intervals of 7 x 2 min, under vacuum).

Samples were then stained with DAPI (20 min, 2 µg/ml in PHEM), rinsed twice in PHEM and kept in the same buffer during confocal acquisition. We identified putative choano-neuroid cells based on the presence of DAPI staining in the center of a choanocyte chamber. Locations of target choanocyte chambers, hosting a central putative neuroid cell, were mapped in three dimensions using the DAPI fluorescence imaged on a Leica SP8 confocal microscope (10x and 20x dry objectives). We acquired images of a large portion of the sponge bodies, focusing on the edges that are the most EM-accessible regions. Confocal acquisition permitted the accumulation of morphological landmarks later used to trace back the selected choanocyte chamber for the FIB-SEM acquisition.

**Post-fixation for FIB-SEM**

After confocal acquisition, samples were post-processed for FIB-SEM using an adaptation of the rOTO protocol (*75*, *76*) aided by microwave processing, as detailed by Schieber et al (*77*). Post-fixation steps are summarized in Table 1. PHEM was prepared by dilution of a 4x PHEM stock, containing PIPES 240 mM, HEPES 100 mM, $MgSO_4$ 8 mM, and

EGTA 40 mM in $ddH_2O$ and adjusted to pH 6.9 with KOH. Durcupan was freshly prepared by mixing the components A-D in the following quantities: A: 11.4 g, B: 10 g, C: 0.3 g and D: 0.1 g.

| | **Post processing steps** | **Duration and incubation mode** |
|---|---|---|
| 1. | 1% $OsO_4$ and 1.5% $K_4Fe(CN)_6$ in PHEM | In microwave, 7 x 2 min at 150 W cycling on and off, vacuum on |
| 2. | Rinse 2 times in PHEM | 3 min each in the microwave |
| 3. | Rinse 5 times in water | 3 min each in the microwave |
| 4. | 1% TCH in water | 10 min on the bench |
| 5. | OsO4 1% in water | In microwave, 7 x 2 min cycling on and off at 150 W, vacuum on |
| 6. | Rinse 2 times in water | 3 min each rinse in the microwave |
| 7. | 1% UA in the microwave | 7 x 1 min at 150 W cycling on and off in vacuum |
| 8. | Dehydration series in ethanol (10%,25%,50%,75%,90%,95%,100%x3) | 40 sec each in the microwave, vacuum off |
| 9. | Infiltration series in Durcupan diluted in ethanol, repeat twice each step: 10%,30%,50%,70%,90%,100% | 2 min each in the microwave, vacuum on |
| 10. | Change to freshly prepared Durcupan resin for flat embedding (by leaving an about 2mm thick resin layer on the coverslip) and polymerize | 72 hrs at 60°C in regular oven |

The glass coverslips were detached from the resin surface using multiple heat shocks in liquid nitrogen and hot water. Resin blocks were then trimmed down to the size of the individual sponges, into 3-4 mm large and about 2 mm thick blocks.

**Targeted trimming for volume EM:**

Targeted trimming is necessary to expose the region of interest to FIB-SEM imaging, i.e. placing the targeted cell not deeper than 10 to 30 µm from the block surface, both in the imaging and in the milling directions. To achieve this step with precision, the topology of the resin embedded sponges was characterized by a combination of transmitted light and microscopic X-ray computed tomography (microCT) imaging (performed with a Bruker SkyScan 1272). Using a 3D CLEM strategy (*78*, *79*) - the confocal data was overlaid to the images from the embedded sample, using Amira 6.5, thus predicting the position of the target choanocyte chamber. With this information the blocks were further trimmed using a glass knife and an ultramicrotome (Leica EM UC7), creating a surface 10 µm above the target. Note that it is recommended not to use diamond knives on this sample, as *Spongilla* produces thick and hard glass spicules. The trimmed blocks were then mounted on FIB-SEM stubs using conductive epoxy resin, polymerised overnight at 60 °C. Finally a layer of silver paint and gold sputter coating were applied.

**Volume EM by FIB-SEM acquisition:**

We imaged the samples on a Zeiss Crossbeam 540, using SmartSEM (Zeiss) to setup the instrument and Atlas 3D v5 to define and guide the acquisition. The surface of the block was platinum-coated in correspondence of the predicted location of the target choanocyte chamber. Acquisition occurred in three separate sessions for a cumulative time of about 96 hours, during which the acquisition area was continuously optimized in order to accommodate the full choanocyte chamber and minimize charging. The final acquired volume was 60 µm x 60 µm x 50 µm with an isotropic pixel resolution of 15 nm.

**FIB-SEM data pre-processing:**

The resulting image series was loaded as a Virtual stack in Fiji, cropped to the regions of interest and converted to 8-bit using batch processing. Alignment occurred in two steps: first by SIFT alignment in Fiji (image J ref) and then using a custom written Python script for subpixel alignment.

**Data visualization and rendering:**

A selection of cells, cilia and microvilli were segmented as detailed above, exported as .obj meshes into Blender 8.0 for visualization and rendering. For visualization of the whole volume the pre-processed image sequence was binned 2 x 2 x 2, corrected for the differences in background between the different acquisition sessions and imported into Drishti v2 (*80*), using the drishtiimport.exe program.

To study the interaction of choancytes and central cells in the EM volume, we produced an instance segmentation of the relevant structures, namely cell bodies, microvilli, and cilia of choanocytes, apopylar cells, and choano-neuroid cells. The size, shape and frequency of objects of these three categories differ significantly. In addition the intensity histogram shifts between parts of the data-set that were imaged in different sessions. These facts make the task at hand too challenging to be solved by a thresholding based approach. Thus, we employed the Lifted Multicut (*81*) based segmentation workflow developed in Pape et al. (*82*) for instance segmentation problems in bio-medical images. This approach consists of two main steps: first a machine learning approach is used to predict per voxel probabilities for object boundaries and semantic classes. Based on these predictions, a graph-based segmentation problem is formulated, where boundary evidence is mapped to local interactions and semantic evidence to non-local interactions.

Here, we use the Autocontext workflow of ilastik (*83*) for the first step. This algorithm consists of multiple stages of voxel classification. In an individual stage a Random Forest Classifier (*84*) predicts semantic class probabilities for each voxel. The classifier is trained from sparse labels with ilastik and uses the stacked responses of several convolutional filters applied to the input data as features. Each stage is presented with the raw data and the predictions of the previous stage; thus refining the previous predictions. We perform three stages of Autocontext:

in the first and second stage, we predict six different classes: *background*, *object boundary* (corresponding to membranes of cells, flagella and microvilli), c*ytoplasm*, *nucleus*, *flagellum* and *microvillus*. In the third stage, we predict the binary classes *boundary* and *non-boundary*.

We use the boundary predictions from the third stage and the class predictions from the second stage to perform Lifted Multicut based segmentation. This approach involves several sub-steps: first, the boundary predictions are used to compute supervoxels via distance-transform watersheds (see *85* for details). These supervoxels are mapped to a graph which represents each supervoxel by a node. Two nodes are connected via an edge if the corresponding supervoxels are adjacent. In order to formulate the segmentation problem, we compute edge weights from the boundary predictions; these weights express the likelihood that the incident nodes belong to the same object. Following the procedure outlined in Pape et al. (*82*), we augment this graph by sparse lifted edges. To derive these edges we first compute the connected component segmentation of the thresholded microvilli and flagella predictions. Then we map the resulting objects to graph nodes via overlap with the corresponding supervoxels and introduce lifted edges between nodes mapped to the same objects. We associate attractive weights with the lifted edges. Based on the graph and lifted edges, we formulate a lifted Multicut graph partitioning problem and solve it with the hierarchical Lifted Multicut solver introduced in Pape et al. (*82*) as an extension of the Multicut solver (*86*).

Note that solving a Multicut problem based on the graph without lifted edges can also yield a segmentation. However, we observe that this segmentation heavily over-segments microvilli and flagella. Both structures have an elongated shape with small diameter, which results in small supervoxels. This results in a non-robust estimate for the boundary evidence based edge weights and causes the over-segmentation. Hence, we add the attractive lifted edges derived from more robust semantic predictions to alleviate the degree of over-segmentation. Also note that the segmentation obtained via connected components on the thresholded predictions on its own is not of sufficient quality, mainly because it falsely merges several cells. As a last step, we manually proofread objects involved in the interactions studied closer using Paintera (https://github.com/saalfeldlab/paintera), and in some cases manually merged the microvilli from a single choanocyte collar into a single object.

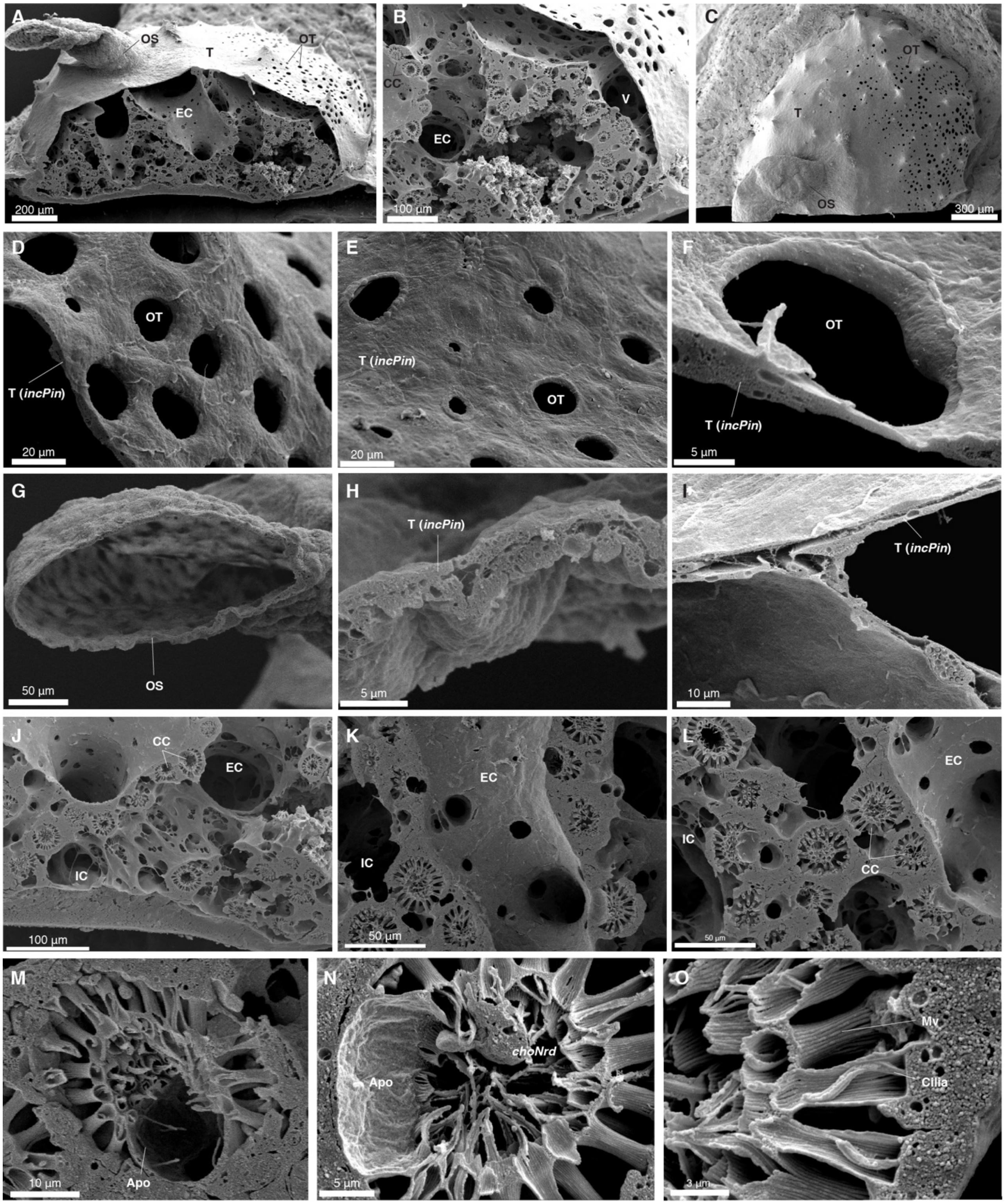

A
OS
T
OT
EC
200 µm
B
CC
V
EC
100 µm
C
OT
T
OS
300 µm
D
OT
T (incPin)
20 µm
E
T (incPin)
OT
20 µm
F
OT
T (incPin)
5 µm
G
OS
50 µm
H
T (incPin)
5 µm
I
T (incPin)
10 µm
J
CC
EC
IC
100 µm
K
EC
IC
50 µm
L
EC
IC
CC
50 µm
M
Apo
10 µm
N
choNrd
Apo
5 µm
O
Mv
Cilia
3 µm

**Figure S1. SEM reveals gross morphology of *Spongilla lacustris*.**

(A-B) Lateral view cross-section of juvenile *S. lacustris*. CC – choanocyte chamber, EC – excurrent canal, OS – osculum, OT – ostia, T – epithelial tent. V – vestibule.

(C) Overhead view.

(D-F) Images of ostia, incurrent opennigs in the outer epithelial tent through which water enters the vestibule.

(G) Images of the osculum, the final excurrent structure through which water exits the sponge.

(H-I) Cross-sections of the outer epithelial tent.

(J-L) Choanocyte chambers embedded in the canal system. IC – incurrent canal.

(M-N) Choanocyte chambers with ciliated apopylar cells forming an excurrent pore. Apo – apopylar pore.

(O) Cross-section of choanocyte microvilli collars and cilia. Mv – microvillus.

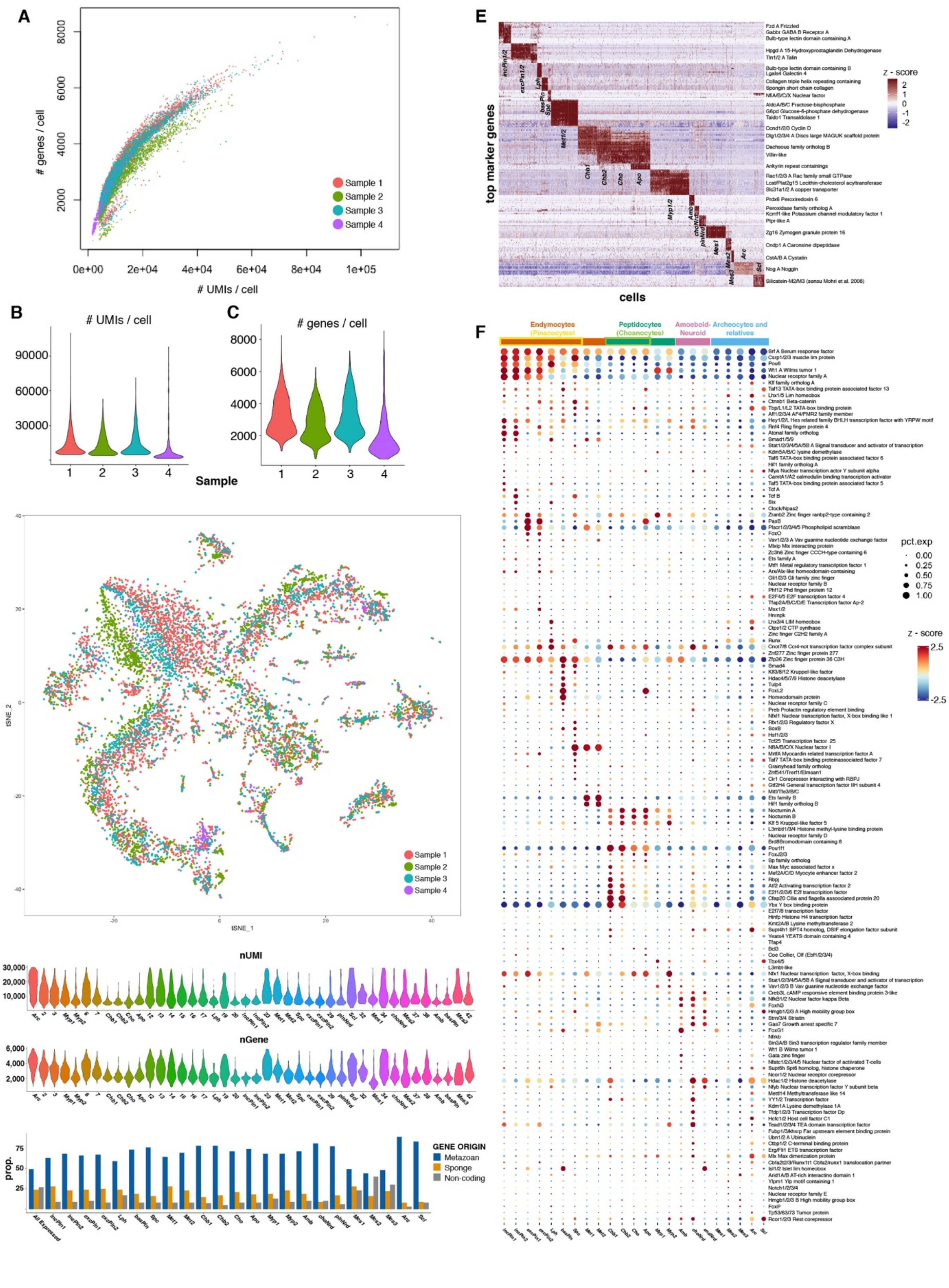
A
### genes / cell
### UMIs / cell
Sample 1
Sample 2
Sample 3
Sample 4
B
### UMIs / cell
C
### genes / cell
Sample
D
tSNE_1
tSNE_2
E
top marker genes
cells
z - score
F
Endymocytes (Pinacocytes)
Peptidocytes (Choanocytes)
Amoeboid-Neuroid
Archeocytes and relatives
pct.exp
z - score
G
nUMI
H
nGene
I
prop.
GENE ORIGIN
Metazoan
Sponge
Non-coding

**Figure S2. Cell type clustering and hierarchical relationships from whole-body scRNAseq of *S. lacustris*.**

(A) UMI count vs. expressed gene count for all cells (colored by sample).

(B) Violin plots of UMI count distribution for each sample.

(C) Violin plots of gene count distribution for each sample.

(D) tSNE plot colored by sample.

(E) Heatmap of top markers for *Spongilla* cell types.

(F) Dotplot showing all expressed transcription factors.

(G) Violin plots of UMI count distribution for each of 42 distinct clusters (cell types and transitional clusters).

(H) Violin plots of gene count distribution for each of 42 distinct clusters (cell types and transitional clusters).

(I) Gene classes for top 100 markers from each cell type. Metazoan – protein-coding gene originating in animal ancestor. Sponge – protein-coding genes originating within Porifera or in poriferan stem line. Non-coding – does not contain at least 70 amino acid open reading frame.

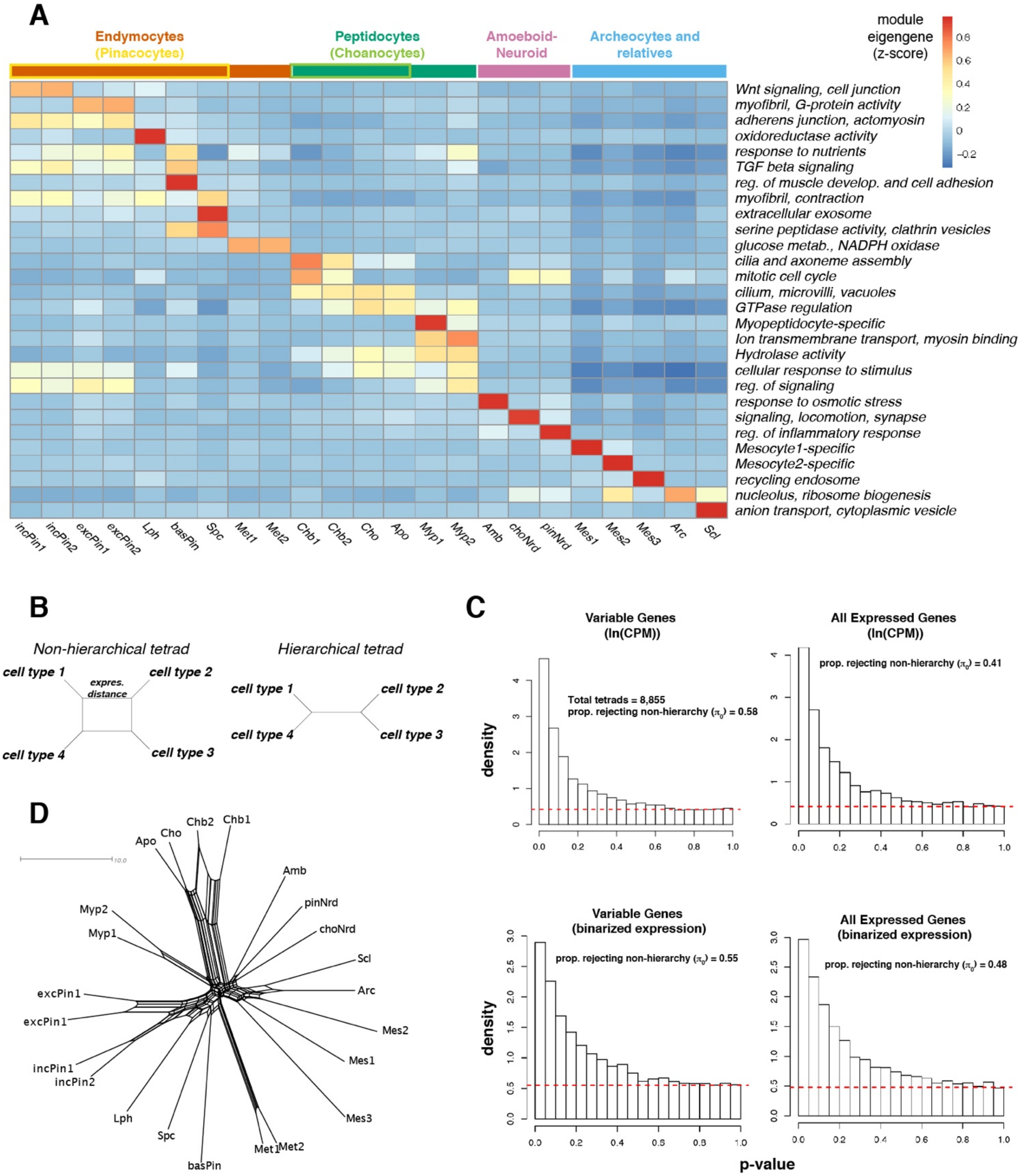
A
Endymocytes
(Pinacocytes)
Peptidocytes
(Choanocytes)
Amoeboid-
Neuroid
Archeocytes and
relatives
module
eigengene
(z-score)
0.8
0.6
0.4
0.2
0
−0.2
Wnt signaling, cell junction
myofibril, G-protein activity
adherens junction, actomyosin
oxidoreductase activity
response to nutrients
TGF beta signaling
reg. of muscle develop. and cell adhesion
myofibril, contraction
extracellular exosome
serine peptidase activity, clathrin vesicles
glucose metab., NADPH oxidase
cilia and axoneme assembly
mitotic cell cycle
cilium, microvilli, vacuoles
GTPase regulation
Myopeptidocyte-specific
Ion transmembrane transport, myosin binding
Hydrolase activity
cellular response to stimulus
reg. of signaling
response to osmotic stress
signaling, locomotion, synapse
reg. of inflammatory response
Mesocyte1-specific
Mesocyte2-specific
recycling endosome
nucleolus, ribosome biogenesis
anion transport, cytoplasmic vesicle
incPin1
incPin2
excPin1
excPin2
Lph
basPin
Spc
Met1
Met2
Chb1
Chb2
Cho
Apo
Myp1
Myp2
Amb
choNrd
pinNrd
Mes1
Mes2
Mes3
Arc
Scl
B
Non-hierarchical tetrad
Hierarchical tetrad
cell type 1
cell type 2
cell type 3
cell type 4
expres. distance
C
Variable Genes
(ln(CPM))
All Expressed Genes
(ln(CPM))
Total tetrads = 8,855
prop. rejecting non-hierarchy ($\pi_0$) = 0.58
prop. rejecting non-hierarchy ($\pi_0$) = 0.41
Variable Genes
(binarized expression)
All Expressed Genes
(binarized expression)
prop. rejecting non-hierarchy ($\pi_0$) = 0.55
prop. rejecting non-hierarchy ($\pi_0$) = 0.48
density
p-value
D
10.0
Apo
Cho
Chb2
Chb1
Amb
pinNrd
choNrd
Scl
Arc
Mes2
Mes1
Mes3
Met1
Met2
basPin
Spc
Lph
incPin1
incPin2
excPin1
excPin1
Myp1
Myp2

**Figure S3. Cell type clustering and hierarchical relationships from whole-body scRNAseq of *S. lacustris*.**

(A) Heatmap showing eigengene expression for all gene models inferred from distinct cell type clusters.

(B) Box plots depicting example non-hierarchical (top) and hierarchical (bottom) relationships of cell type tetrads.

(C) P-value distributions from treeness tests of 8,855 cell type tetrads. Red dashed line indicates uniform distribution of p-values expected under null hypothesis.

(D) SplitsTree network of *S. lacustris* cell type relationships.

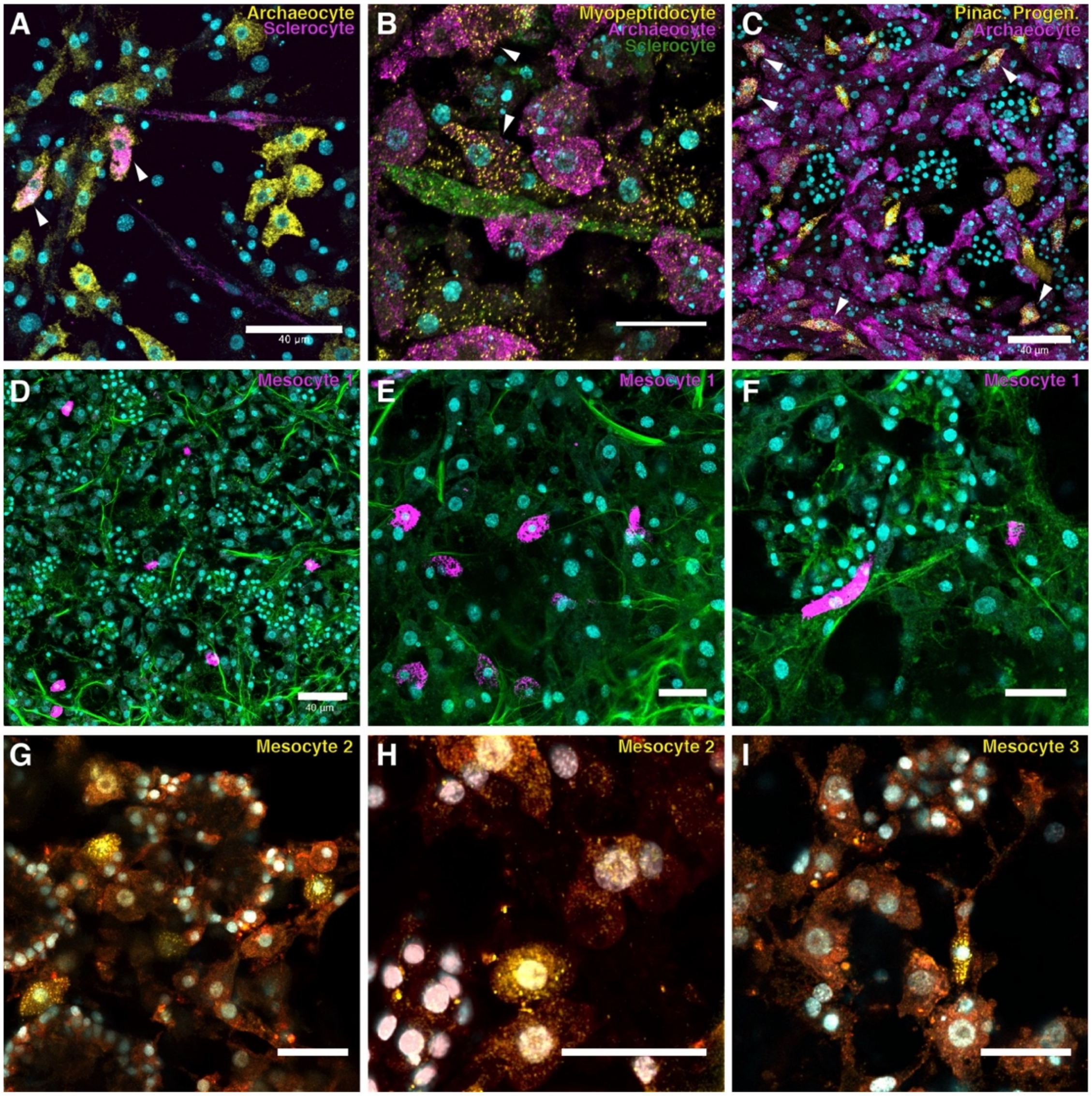


**Figure S4. smFISH of cell types in the archaeocyte and relatives family.**

(A-C) *Eef1a1* labels archaeocytes with prominent nucleolus as well as cells co-staining cells with markers for distinct cell types, including sclerocytes (*Silicatein M2-M3*), myopeptidocytes (*Hpgds*), and pinacocyte progenitors (*c102743_g2*). Scale bars 20µm except where noted. Membrane stains are CellBrite Fix (green) or Fm143-Fx (red). CellBrite Fix also labels intercellular fibrils.

(D-F) Mesocyte 1 marker (c102051_g1) stains medium-to-large cells scattered throughout sponge mesohyl.

(G-H) Mesocyte 2 marker (ortholog of Drosophila *Dip-B*) identifies medium-sized ovoid-shaped cells in the mesohyl.

(I) Mesocyte 3 marker (c102181_g1) stains very rare mesenchymal cells that are fusiform in shape.

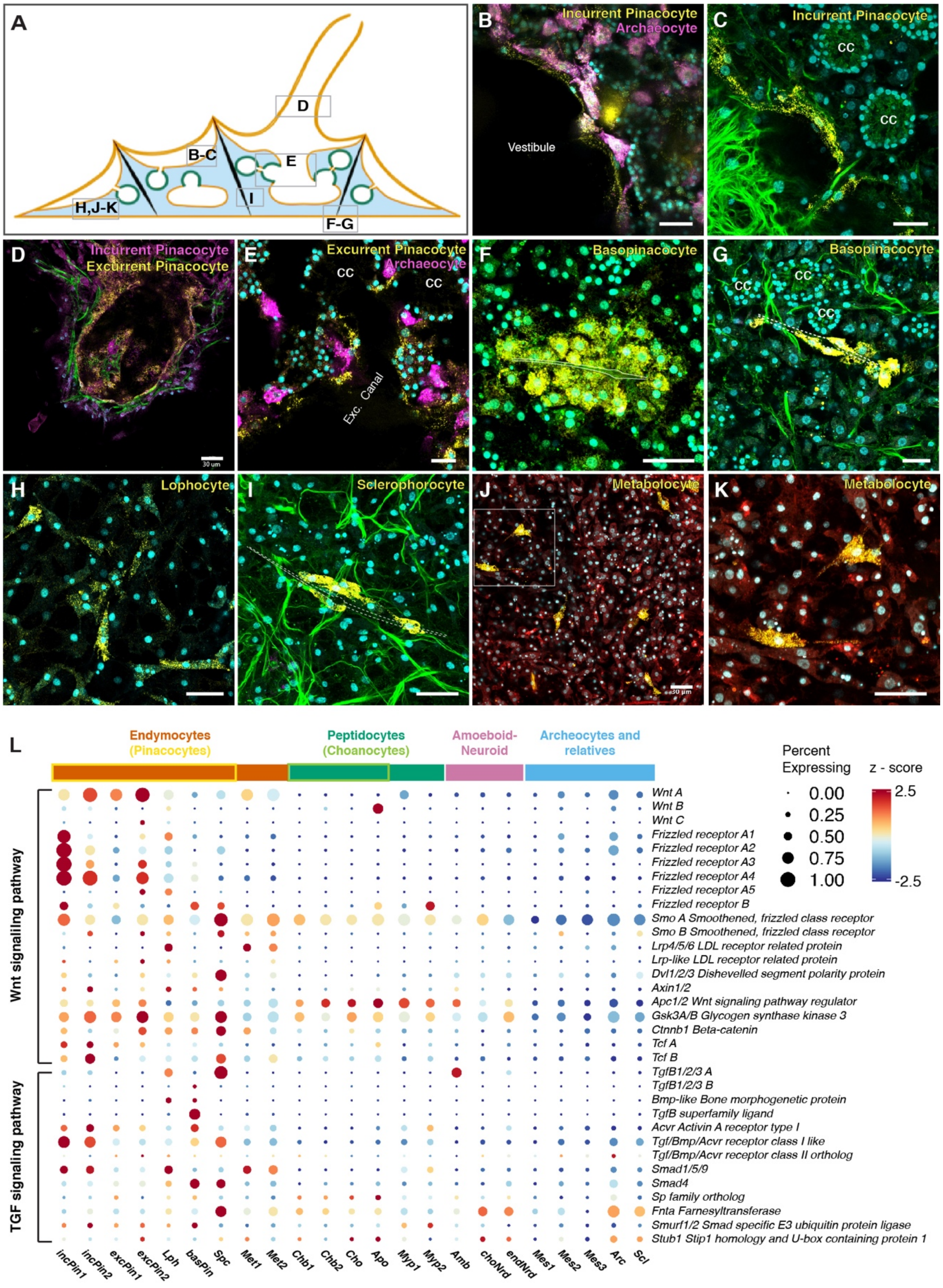
A
D
B-C
E
H,J-K
I
F-G
B
Incurrent Pinacocyte
Archaeocyte
Vestibule
C
Incurrent Pinacocyte
cc
D
Incurrent Pinacocyte
Excurrent Pinacocyte
30 μm
E
Excurrent Pinacocyte
Archaeocyte
cc
Exc. Canal
F
Basopinacocyte
G
Basopinacocyte
cc
H
Lophocyte
I
Sclerophorocyte
J
Metabolocyte
K
Metabolocyte
L
Endymocytes
(Pinacocytes)
Peptidocytes
(Choanocytes)
Amoeboid-
Neuroid
Archeocytes and
relatives
Percent
Expressing
0.00
0.25
0.50
0.75
1.00
z - score
2.5
-2.5
Wnt signalling pathway
TGF signaling pathway
Wnt A
Wnt B
Wnt C
Frizzled receptor A1
Frizzled receptor A2
Frizzled receptor A3
Frizzled receptor A4
Frizzled receptor A5
Frizzled receptor B
Smo A Smoothened, frizzled class receptor
Smo B Smoothened, frizzled class receptor
Lrp4/5/6 LDL receptor related protein
Lrp-like LDL receptor related protein
Dvl1/2/3 Dishevelled segment polarity protein
Axin1/2
Apc1/2 Wnt signaling pathway regulator
Gsk3A/B Glycogen synthase kinase 3
Ctnnb1 Beta-catenin
Tcf A
Tcf B
TgfB1/2/3 A
TgfB1/2/3 B
Bmp-like Bone morphogenetic protein
TgfB superfamily ligand
Acvr Activin A receptor type I
Tgf/Bmp/Acvr receptor class I like
Tgf/Bmp/Acvr receptor class II ortholog
Smad1/5/9
Smad4
Sp family ortholog
Fnta Farnesyltransferase
Smurf1/2 Smad specific E3 ubiquitin protein ligase
Stub1 Stip1 homology and U-box containing protein 1
incPin1
incPin2
excPin1
excPin2
Lph
basPin
Spc
Met1
Met2
Chb1
Chb2
Cho
Apo
Myp1
Myp2
Amb
choNrd
endNrd
Mes1
Mes2
Mes3
Arc
Scl

**Figure S5. Endymocyte family.**

(A) Diagram of juvenile sponge illustrating the location of smFISH panels.

(B-K) smFISH of endymocyte cell type markers. Markers: (B) *Tropomyosin* and *Eef1a1*, (C) *Fibcd1 A*, (D) *Tropomyosin* and *c99740_g1*, (E) *Tetraspanin* and *Eef1a1*, (F-G) *Spongin short-chain collagen*, (H) *c100322_g1*, (I) *c101435_g1*, (J-K) *Reelin B*. All scale bars 20µm except where noted in the panel. Membrane stains are CellBrite Fix (green) or Fm143-Fx (red). Dotted lines show the outline of the spicule. CC – choanocyte chamber.

(L) Dotplot of Wnt and Tgf signaling pathway genes in *S. lacustris.*

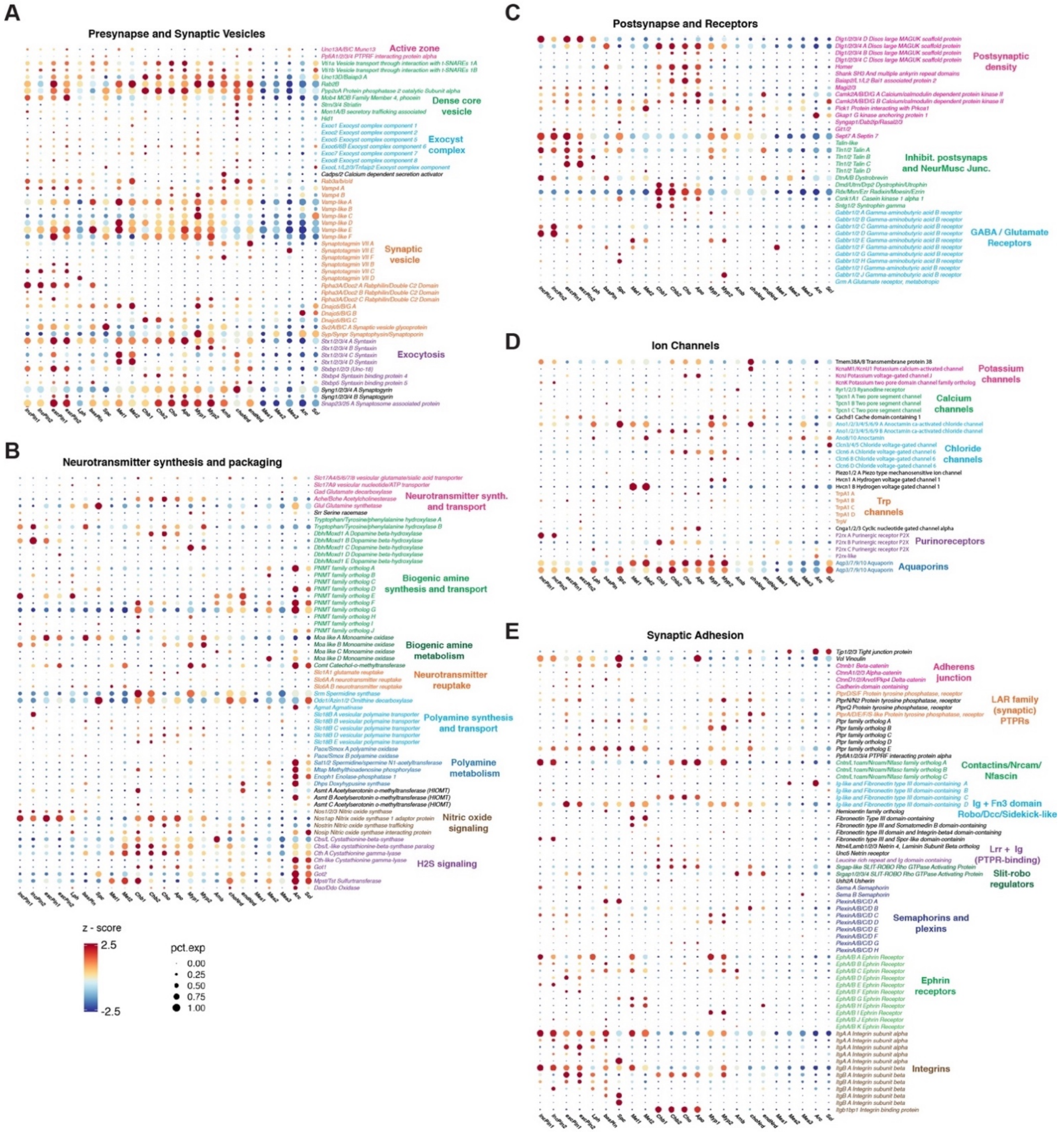


**Figure S6. Neuronal gene expression in *S. lacustris* cell types.**

Dotplots showing relative expression of (A) presynaptic, (B) neurotransmitter synthesis and transport, (C) postsynaptic, (D) ion channel, and (E) synaptic adhesion genes.

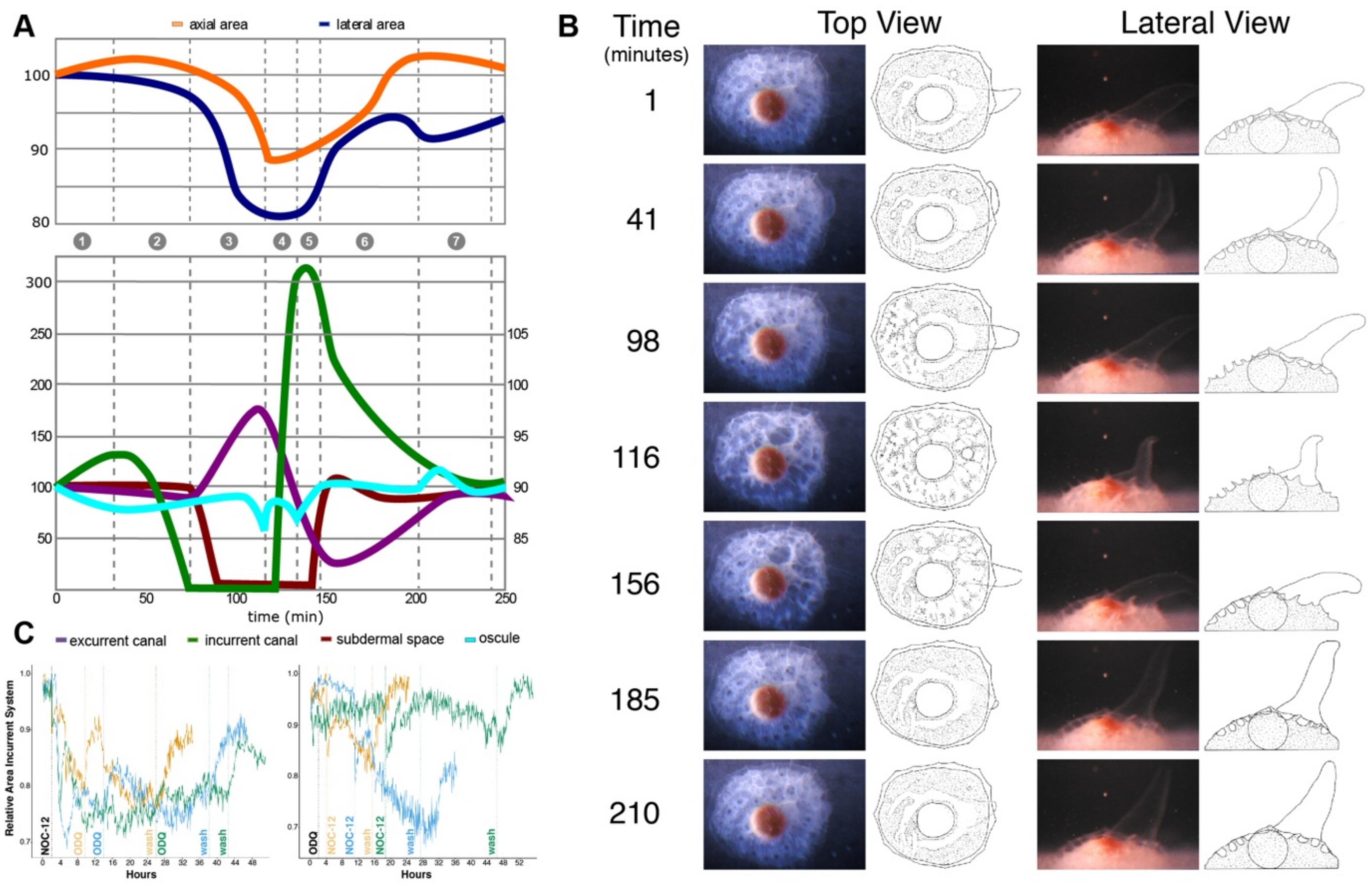


**Figure S7. Contractions in juvenile *S. lacustris*.**

(A) Graph showing changes in the relative area of canal system components during 7 stages of a typical contraction cycle. Colored lines depict relative area of different regions of the canal system.

(B) Brightfield and schematic views of sponge for each of 7 stages of typical contraction cycle. Times (in minutes) correspond to those in the graph in panel A.

(C) Timelapse plot of contraction state following treatment with NOC-12 (C) and ODQ (D) in series. treatment. Colored lines indicate experiments with different periods between successive treatments.

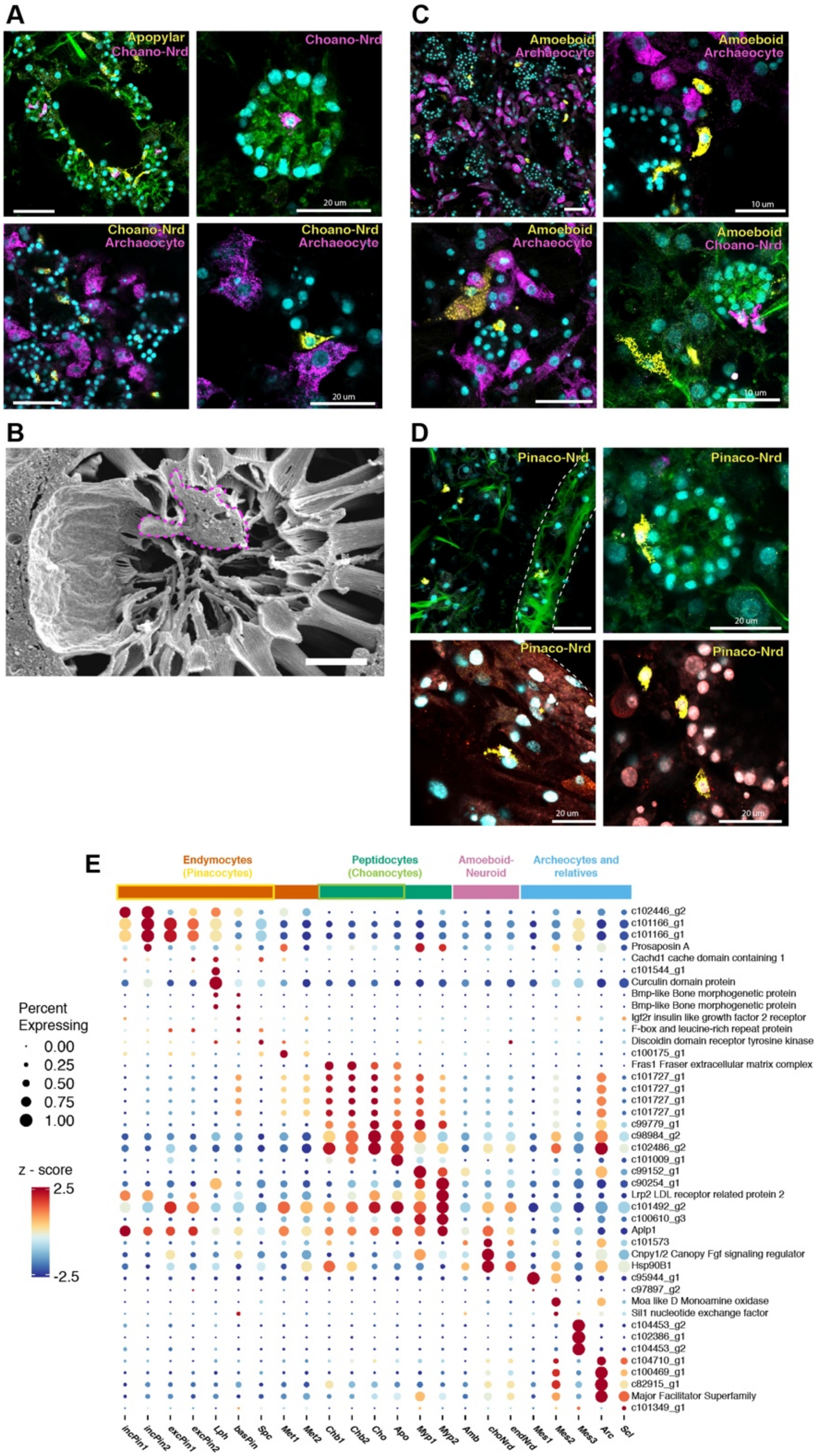
A
Apopylar
Choano-Nrd
Choano-Nrd
20 um
Choano-Nrd
Archaeocyte
Choano-Nrd
Archaeocyte
20 um
B
C
Amoeboid
Archaeocyte
Amoeboid
Archaeocyte
10 um
Amoeboid
Archaeocyte
Amoeboid
Choano-Nrd
10 um
D
Pinaco-Nrd
Pinaco-Nrd
20 um
Pinaco-Nrd
20 um
Pinaco-Nrd
20 um
E
Endymocytes
(Pinacocytes)
Peptidocytes
(Choanocytes)
Amoeboid-
Neuroid
Archeocytes and
relatives
Percent
Expressing
0.00
0.25
0.50
0.75
1.00
z - score
2.5
-2.5
c102446_g2
c101166_g1
c101166_g1
Prosaposin A
Cachd1 cache domain containing 1
c101544_g1
Curculin domain protein
Bmp-like Bone morphogenetic protein
Bmp-like Bone morphogenetic protein
Igf2r insulin like growth factor 2 receptor
F-box and leucine-rich repeat protein
Discoidin domain receptor tyrosine kinase
c100175_g1
Fras1 Fraser extracellular matrix complex
c101727_g1
c101727_g1
c101727_g1
c101727_g1
c99779_g1
c98984_g2
c102486_g2
c101009_g1
c99152_g1
c90254_g1
Lrp2 LDL receptor related protein 2
c101492_g2
c100610_g3
Aplp1
c101573
Cnpy1/2 Canopy Fgf signaling regulator
Hsp90B1
c95944_g1
c97897_g2
Moa like D Monoamine oxidase
Sil1 nucleotide exchange factor
c104453_g2
c102386_g1
c104453_g2
c104710_g1
c100469_g1
c82915_g1
Major Facilitator Superfamily
c101349_g1
incPin1
incPin2
excPin1
excPin2
Lph
basPin
Spc
Met1
Met2
Chb1
Chb2
Cho
Apo
Myp1
Myp2
Amb
choNrd
endNrd
Mes1
Mes2
Mes3
Arc
Scl

**Figure S8. Neuroid-amoeboid Family.**

(A) smFISH of choano-neuroid cells (*Peroxidase*), with archaeocytes (*Eef1a1*) and apopylar cells (*c85989_g1*).

(B) SEM of a putative choano-neuroid cell (outlined by violet dashed line) in choanocyte chamber.

(C) smFISH of amoeboid cells (*c101448_g1* and *c103466_g1*) with archaeocytes (*Eef1a1*) and choano-neuroid cells (*Perodixase*).

(D) smFISH of pinaco-neuroid cells (marker *Acp5*).

(E) Dotplot of putative secreted peptides.

| Cell Type Name | Nomen. | Cell Type Family | Cluster # | Description |
|---|---|---|---|---|
| Archaeocytes | *Arc* | Archaeocyte Family | 1 | Large mesenchymal cells with prominent nucleolus found throughout mesohyl |
| Sclerocytes | *Scl* | Archaeocyte Family | 31 | Elongate mesenchymal cells, usually enwrapping growing spicule |
| Mesocytes 1 | *Mes1* | Archaeocyte Family | 33 | Medium-sized and irregularly-shaped mesenchymal cell lacking nucleolus, fairly common in mesohyl. |
| Mesocytes 2 | *Mes2* | Archaeocyte Family | 36 | Medium-sized egg-shaped mesenchymal cells with nucleolus, smaller than archaeocytes |
| Mesocytes 3 | *Mes3* | Archaeocyte Family | 41 | Elongate mesohyl cell, rare |
| Incurrent Pinacocytes1/2 | *incPin1/2* | Endymocytes | 21-22 | Epithelial cells forming external tent, outer layer of osculum, and lining subdermal lacunae |
| Excurrent Pinacocytes1/2 | *excPin1/2* | Endymocytes | 27-28 | Epithelial cells lining excurrent channels and inner layer of osculum |
| Basopinacocytes | *basPIN* | Endymocytes | 40 | Epithelial cells forming layer adherent to substratum. Also enwrap spicules inserted into the base of sponge. |
| Sclerophorocytes | *Spc* | Endymocytes | 26 | Cells typically observed enwrapping mature spicules, may play role in spicule transport |
| Lophocytes | *Lph* | Endymocytes | 18 | Migratory cells with elongate, asymmetric shape. Most prevalent near basopinacocytes and incurrent pinacocytes. |
| Metabaloyctes1/2 | *Met1/2* | Endymocytes | 24-25 | Large, multipolar mesohyl cells lacking nucleolus. |
| Choanoblasts1/2 | *Chb1/2* | Peptidocytes | 8-9 | Proliferative medium-size cells typically adjacent to choanocyte chambers, lacking collar and cilium |
| Choanocytes | *Cho* | Peptidocytes | 10 | Mature choanocytes, small cells with collar and cilium |
| Apopylar Cells | *Apo* | Peptidocytes | 11 | Cells forming the excurrent pore of mature choanocyte chamber. Cilia present but lacks collar. |
| Myopeptidocytes1/2 | *Myp1/2* | Peptidocytes | 4-5 | Medium-sized mesechymal cells with prominent vacuoles and long projections forming cellular network; nucleolus absent |
| Amoeboid | *Amb* | Amoeboid-Neuroid | 39 | Small mesenchymal cells, sometimes observed engulfing other cells |
| Neuroid (choanocytes) | *choNrd* | Amoeboid-Neuroid | 35 | Small, multipolar mesenchymal cells with thin projections, usually found in choanocyte chambers |
| Neuroid (pinacocytes) | *pinNrd* | Amoeboid-Neuroid | 30 | Small, multipolar mesenchymal cells with thin projections, scattered in mesenchyme and in proximity to pinacocytes |

**Table S1. Cell types in *Spongilla lacustris*, their morphology and distribution.**

**Movie S1.** Simultaneous top-down and lateral views of endogenous sponge contractions in juvenile *S. lacustris*.

**Movie S2.** Noc-12 treatment of juvenile *S. lacustris*.

**Movie S3.** ODQ treatment of juvenile *S. lacustris*.

**Movie S4.** Rendering of FIB-SEM data reveals choano-neuroid cells in choanocyte chamber.

**Movie S5.** Rendering of segmented FIB-SEM data illustrates interactions between choano-neuroid cells and individual choanocytes.

**Data S1.** Cell type marker genes and smFISH oligo sequences.

**Data S2.** Klf family gene phylogenies.

**Data S3.** Homeobox family gene phylogeny.
