## Supplementary figures and images for "Profiling cellular diversity in sponges informs animal cell type and nervous system evolution"

### Data S2

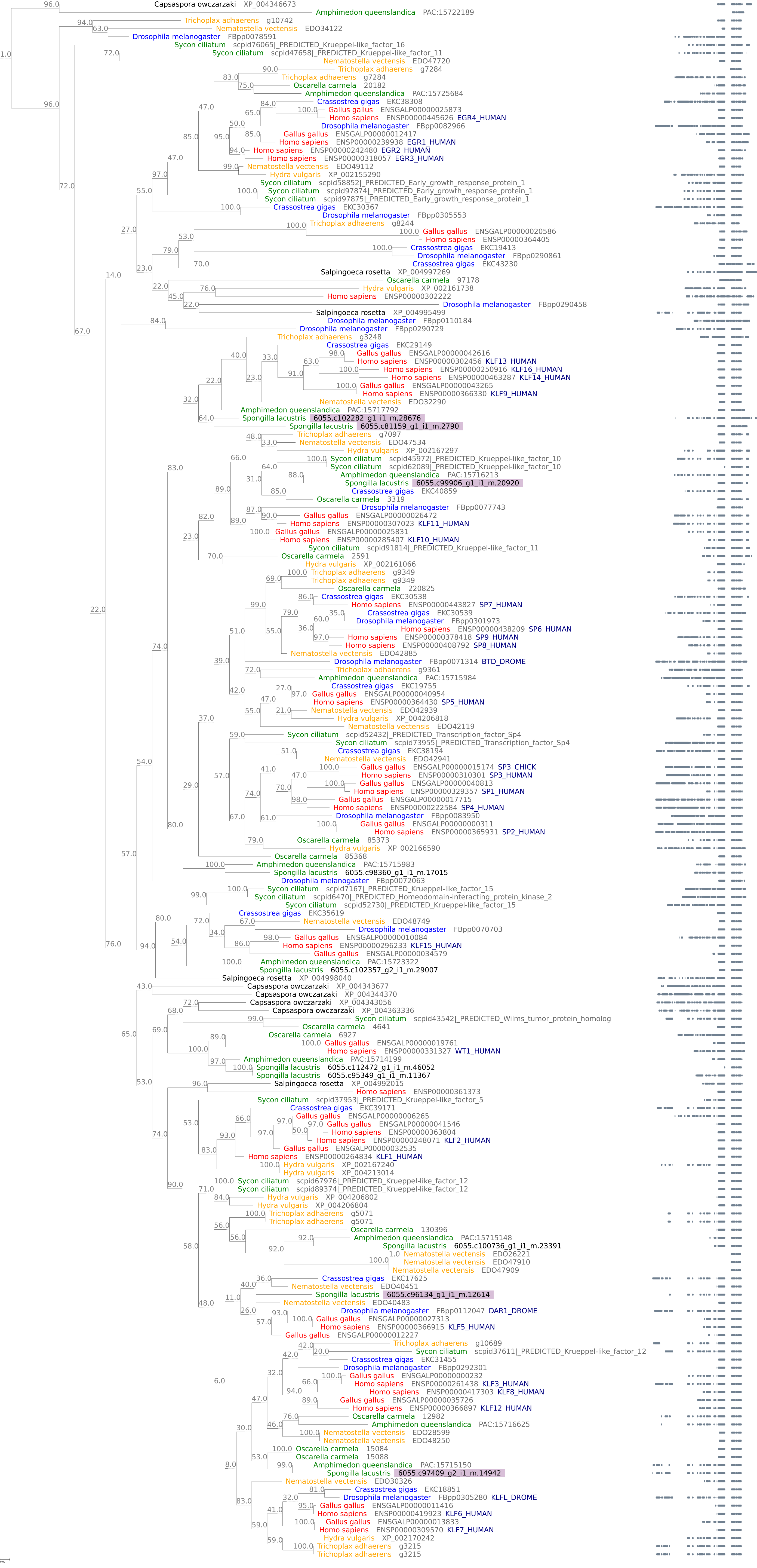

### Data S3

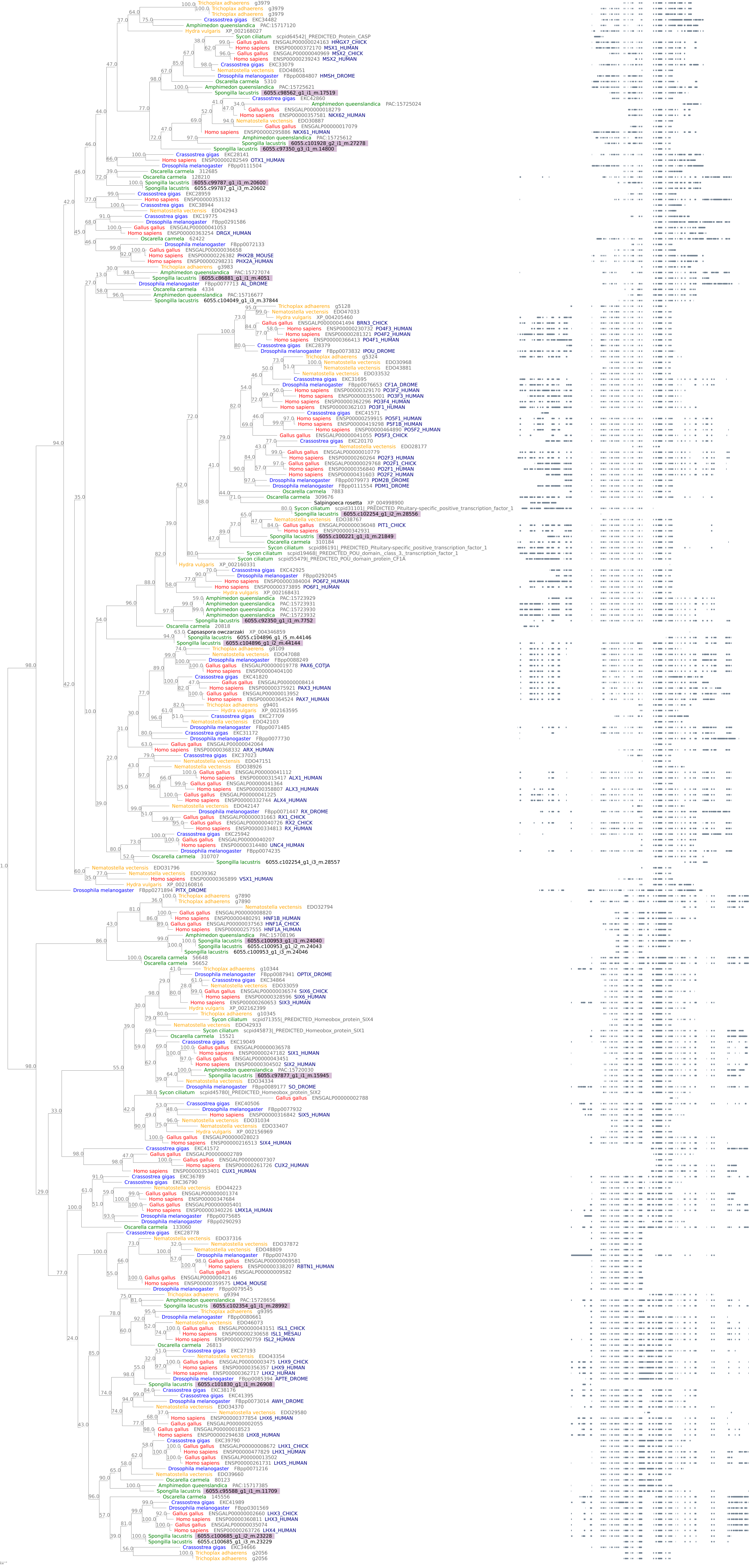
